# Recruitment limitation increases susceptibility to fishing-induced collapse in a spawning aggregation fishery

**DOI:** 10.1101/2023.10.16.562228

**Authors:** Erica T. Jarvis Mason, Thomas V. Riecke, Lyall F. Bellquist, Daniel J. Pondella, Brice X. Semmens

**Affiliations:** Scripps Institution of Oceanography, University of California San Diego, California, USA; Department of Ecosystem and Conservation Sciences, University of Montana, USA; California Oceans Program, The Nature Conservancy, California, USA; Vantuna Research Group, Occidental College, California, USA

## Abstract

Aggregation-based fisheries are notorious for booms and busts driven by aggregation discovery and subsequent fishing-induced collapse. However, environment-driven sporadic recruitment in some since-protected populations has delayed recovery, suggesting recruitment-limitation may be a key driver of their population dynamics and fishery recovery potential. To glean insight into this dynamic, we focused on an overexploited temperate aggregate spawner (Barred Sand Bass; *Paralabrax nebulifer*) and leveraged a long-term mark-recapture data set spanning different oceanographic and harvest histories in a custom Bayesian capture-mark-reencounter modeling framework. We coupled this demographic analysis with long-term trends in sea surface temperature, harvest, adult and juvenile densities, and historical accounts in the literature. Our results point to a history of multidecadal windows of fishing opportunity and fishing-induced collapse that were largely driven by sporadic, warm water recruitment events, which may be externally sourced. Nevertheless, we found that environment-driven sporadic recruitment was not a factor impeding recovery following the last collapse, as recruitment remained elevated due to novel, anomalously warm conditions. Despite signs of incipient population recovery, spawning aggregations remain absent, indicating other potential factors (e.g., continued fishing during spawning season, residual Allee effects) have delayed fishery recovery to date. Aggregate spawner populations that are dependent on sporadic recruitment, especially those at their geographic margins, are thus highly susceptible to sudden and potentially extended periods of collapse, making them ill-suited to high CPUE fishing that occurs on spawning grounds. If the goal is to balance the protection of spawning aggregations with long-term fishery sustainability, then limiting aggregation-based fishing during spawning season may be the best insurance policy against collapse and recovery failure.

## 1. INTRODUCTION

Fishes that form large spawning aggregations (i.e., aggregate spawners) are commonly exploited by artisanal, recreational, and commercial fisheries worldwide. However, they are highly vulnerable to overfishing due to the spatiotemporal predictability of their aggregations and other life-history characteristics typical of aggregate spawners (e.g., slow growth, depensatory dynamics, Sadovy de Mitcheson 2016). This is especially true for transient aggregate spawners that migrate long distances to form exceptionally large spawning aggregations for weeks to months (Domeier & Colin 1997, Heyman 2019). Indeed, overfishing has contributed to the collapse of many fisheries based on transient spawning aggregations (Chollett et al. 2020), and recovery has taken decades (Sadovy & Eklund 1999, Aguilar-Perera 2006, Waterhouse et al. 2020) or failed to occur altogether (Perälä et al. 2022).

The delay or lack of recovery in overfished aggregate spawner populations despite measures to enhance populations, is contrary to compensatory population dynamics, in which traditional fisheries management is rooted (e.g., the assumption that *per capita* population growth rate increases at low stock sizes). One explanation includes the Allee effect (Allee 1931, 1938, Stephens et al. 1999), also referred to as depensation in fisheries science, in which a population’s *per capita* growth rate declines upon reaching a low level (i.e., the “Allee-effect threshold”, Hutchings 2015). In an extreme case, a population could potentially be fished to a point at which densities are so low that it is unable to replenish itself (i.e., reaching yet another threshold, “the Allee threshold”, Hutchings 2015), but this can be difficult to detect unless stock sizes are reduced to <1% of unfished spawning biomass (Liermann and Hilborn 2001, Hilborn et al. 2014). It is also possible that residual Allee effects could delay fishery recovery even after a population begins to rebound. For example, in aggregation-based fisheries, low densities of adults could potentially disrupt the behavioral dynamics of spawning aggregation formation, resulting in the loss of generational transfer of historical spawning ground locations (Warner 1988, 1990, Bolden 2000, Semmens et al. 2008).

Yet another factor that could delay recovery in aggregation-based fisheries is recruitment limitation, or environment-driven sporadic recruitment (Semmens et al. 2007, Stock et al. 2021). Although there is evidence that recovery is possible when fishing mortality is majorly curtailed (Hilborn et al. 2014, Chollett et al. 2020, Waterhouse et al. 2020), such recoveries are subject to environmental drivers, with many fished populations showing recruitment fluctuations driven by oceanography (i.e., transport, temperature regimes) rather than (or in addition to) spawning stock biomass (Vert-Pre et al. 2013, Szuwalski et al. 2015). Such variability in recruitment mediates both the resilience of the stock to overfishing, and the determinism of stock recovery following management (Kuparinen et al. 2014). For example, a population may have experienced recruitment-limitation prior to discovery of the fishery and aggregation-based fishing on spawning grounds would have acted to further limit recruitment potential during periods of unfavorable conditions, accelerating the imminent “bust” trajectory and delaying recovery.

One prominent case of delayed recovery in a transient aggregate spawner fishery is that of Barred Sand Bass (Family Serranidae, *Paralabrax nebulifer*) in southern California, USA. Barred Sand Bass (hereafter, BSB) were once the target of a highly popular aggregation-based *recreational-only* fishery during the summer spawning season (Love et al. 1996a, Erisman et al. 2011, Jarvis et al. 2014). Historically, BSB would migrate on average tens of kilometers to form massive spawning aggregations at several locations along the coast (Jarvis et al. 2010, Teesdale et al. 2015), in which the spawning grounds became well-known BSB fishing “hot spots” (Love et al. 1996a). During the 1980s and 1990s, BSB fishing was a focal summer pastime, but a sharp decline in catch-per-unit-effort (CPUE) in the mid-2000s called into question the sustainability of the fishery (Jarvis et al. 2010, Erisman et al. 2011) and prompted the implementation of tighter fishing regulations in 2013 (i.e., increased minimum size limit and decreased daily bag limit, Jarvis et al. 2014). In the decade since, BSB recreational landings and CPUE have fallen to all-time lows (CDFW 2020), and the spawning aggregations have seemingly disappeared, with no signs of fishery recovery (Bellquist et al. 2017). Adding to the uncertainty of recovery is a lack of population estimates and knowledge of the oceanographic drivers influencing BSB population dynamics. Given the lack of formal stock-recruit data for this fishery, which greatly limits detection of an Allee effect, it seems reasonable to determine whether environment-driven recruitment-limitation can be ruled out as a driver of population dynamics and delayed recovery in this fishery.

Though the effect of fishing on the decline of the BSB fishery is well documented (Erisman et al. 2011, Jarvis et al. 2014, Miller & Erisman 2014, Bellquist et al. 2017), the contribution of changing ocean conditions to this decline and lack of recovery remains poorly understood. Temporal trends in fishery-independent data suggest BSB larval and juvenile recruitment in southern California has fluctuated in response to environmental conditions (Stephens et al. 1986, 1994, Jarvis et al. 2014), generally favoring warmer oceanographic climates (Moser et al. 2001, Hsieh et al. 2005, Jarvis et al. 2014). BSB are commonly distributed from Bahia Magdalena in southern Baja California, Mexico to Santa Cruz, CA, USA (Heemstra 1995, Love & Passerelli 2020), but they are rare north of Pt. Conception. Historically, their distribution and availability in California was considered tightly coupled to warm water conditions (Hubbs 1948, Young 1969, Frey 1971, Feder et al. 1974). If BSB recruitment in southern California is more closely tied to environmental conditions than spawning stock biomass, it is likely that climate change will drive changes in recruitment frequency/intensity, in addition to a shift in the geographic range of the population (Hubbs 1948, Pinsky et al. 2020, Walker et al. 2020a). While predicting future stock status may be challenging, examining the historical population dynamics of the species in southern California, in relation to both harvest and the environment, will likely provide context for the anticipated changes a warming ocean will bring.

Population variability in BSB may be at least partially driven by changes in the cumulative effects of environmental drivers and fishing pressure on mortality. In fished populations, total mortality is the sum of mortality due to fishing and natural causes (i.e., predation, disease, etc.). Fishing mortality may be derived from a formal stock assessment (data-rich fisheries), catch-curve analysis (data-poor fisheries), and/or mark-recapture models, with the latter being the recommended method because it provides direct estimates of total mortality (Pine et al. 2003). Mark recapture models can also estimate the discrete form of fishing mortality (i.e., exploitation or harvest rate), which represents the fraction of the population removed due to fishing. Hence, along with estimates of total harvest in the fishery, one can derive the population size from which harvest was drawn.

Here, we take advantage of long-term mark-recapture data spanning different oceanographic regimes and harvest histories and develop a novel Bayesian capture-mark-reencounter (CMR) framework to glean insight into the long-term population dynamics of BSB. Specifically, we estimate demographic rates (e.g., growth, survival, exploitation) and population size during these regimes and compare them to long-term trends in SST, fishery-independent surveys of adult densities, and harvest. Additionally, we model young-of-the-year densities (juvenile recruitment) as a function of SST, adult densities, and harvest, and look for signs of population recovery in adult and juvenile densities in the last decade. Finally, we attempt to reconcile our findings with historical accounts of BSB distribution and availability in the literature. In doing so, we seek to resolve long standing uncertainty in the role of environment-driven sporadic recruitment in the dynamics of this economically and culturally important aggregation-based fishery.

## 2. MATERIALS AND METHODS

### 2.1. Tagging Studies

We analyzed BSB tagging data collected by the California Department of Fish and Wildlife (CDFW, formerly the California Department of Fish Game) between 1962 and 1970 (1960s) and between 1989 and 1999 (1990s), as well as tagging data collected by researchers at Scripps Institution of Oceanography (SIO), UC San Diego, between October 2012 and February 2015 (2010s). In all three periods, BSB were captured by hook-and-line (a small subset in the 1990s were trawl-caught), measured to the nearest mm TL, tagged with external t-bar tags printed with a unique identification number, “Reward”, and phone number, and subsequently released (see Jarvis et al. 2010 for a detailed description of the CDFW tagging studies). Tagging rewards across study periods included low value non-monetary and monetary incentives (e.g., hats, $5 cash, gas cards).

During the 1960s and 1990s, tagging effort was focused primarily during peak spawning (June-August) and was distributed throughout the southern California coast, including spawning and non-spawning grounds (Jarvis et al. 2010, Fig. 1). In the 2010s, tagging occurred year-round at spawning and non-spawning grounds primarily off San Diego, CA, USA (Fig. 1). Some tagging occurred inside a marine protected area (MPA), in which take is prohibited year-round. We filtered the 2010s data to exclude fish tagged in the MPA, as it is likely these fish had a lower probability of capture by anglers restricted to fishing outside of the MPA.

**Figure 1.**
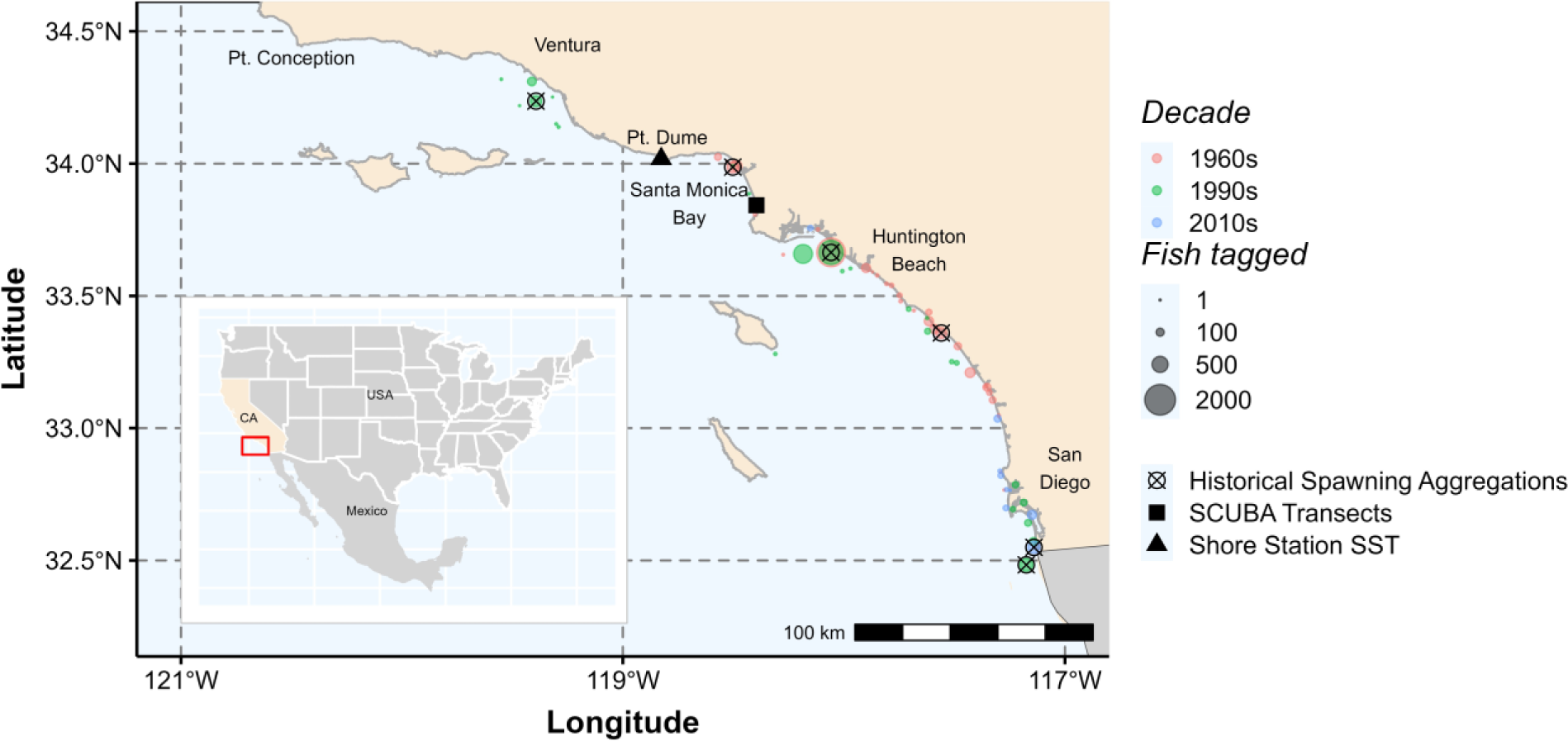
Map of Barred Sand Bass tagging locations (colored circles) in southern California (CA), USA, by decade, including the location of SCUBA transects where Barred Sand Bass juvenile and adult density data were collected (solid black square), Shore Station SST (sea surface temperature, solid black triangle) data collection, and historical spawning aggregations (crosshair).

Data processing and filtering of the tag and recapture data (Supplement S1) resulted in capture histories for 6,473 tagged BSB across the three tagging periods (1960s, 1990s, and 2010s), which represented nearly 25 years of data spanning five decades (Fig. 1, Table 1). Both the number of tagged fish and proportion of recaptures was highest in the 1960s and lowest in the 2010s, and the average size of fish tagged increased over time (Table 1).

**Table 1.** Summary tag and recapture statistics by tagging period (1960s: 1962-1970, 1990s: 1989-1999, 2010s: 2012-2015) for Barred Sand Bass tagged and released in southern California, USA.

|  | 1960s | 1990s | 2010s |
| --- | --- | --- | --- |
| No. of years | 9 | 11 | 2.3 |
| Tagged fish<sup>a</sup> | 3,335 | 2,696 | 442 |
| Total length (mm) |  |  |  |
| range | 218 - 551 | 178 - 647 | 171 - 544 |
| mean (± SD) | 303 ± 34 | 339 ± 62 | 386 ± 60 |
| % mature<sup>b</sup> | 89% | 94% | 97% |
| % legal size<sup>c</sup> | 38% | 69% | 75% |
| Recaptured fish | 255 | 130 | 12 |
| Recapture rate | 8% | 5% | 3% |
| Total length (mm)<sup>d</sup> |  |  |  |
| no. reported lengths | 243 | 65 | 1 |
| range | 260 - 577 | 305 - 508 | 375 - 375 |
| mean (± SD) | 330 ± 36 | 348 ± 35 | 375 |
| % mature<sup>b</sup> | 98% | 100% | 100% |
| % legal size<sup>c</sup> | 84% | 100% | 100% |

<sup>a</sup>Fish tagged in the last year during the 1960s and 1990s were not included in the analysis. For the 1960s and 1990s, we include only fish tagged in June-August, and for the 2010s, we include fish tagged monthly from October 2012 through January 2015.
<sup>b</sup>Based on size at 100% maturity ( $\geq 270$ mm).
<sup>c</sup>Legal size during the 1960s and 1990s was $\geq 305$ mm; legal size increased to $\geq 356$ mm in March 2013.
<sup>d</sup>Lengths not reported for all recaptured fish.

### 2.2. Capture-mark-reencounter (CMR) Model

#### 2.2.1. Parameters (Demographic Rates)

We used a Bayesian multistate framework (Kéry & Schaub 2012) to estimate the following four probabilities:

*ϕ_t_* (true survival) = the probability a fish alive at occasion *t* is alive at occasion *t* + 1,
*p_t_* (recapture probability) = the probability a fish at risk of capture at occasion *t* is recaptured by a biologist at occasion *t*,
*κ_t_* (recovery probability or harvest rate/fishing mortality) = the probability a fish is caught and kept by an angler from occasion *t* to *t* +1 and the tag reported (caught and kept and reported at any time from occasion *t* through the interval between *t* and *t* + 1), and
*R_t_* (resighting probability or catch-and-release (CAR) rate) = the probability a fish at risk of capture at occasion *t* is caught and released (i.e., resighted) by an angler in occasion *t* and the tag reported.

Our approach differs from traditional CMR models (Barker 1997, Riecke et al. 2021) in two ways. First, we assumed no permanent emigration and thus, excluded the fidelity parameters (*F* and *F’*). This assumption was based on the CMR area being sufficiently large to include the geographic area for tagging and variability in BSB mean home range size and migration distance to spawning grounds (Mason & Lowe 2010, Jarvis et al. 2010). Second, we excluded any CAR encounters in the interval between survey occasions (1960s: *n* = 47 [18.4%]; 1990s: *n* = 22 [16.9%]; 2010s: *n* = 2 [16.7%]). The Barker model estimates the probability of CAR at *t* + 1 (the non-survey interval), given the fish survives to occasion *t* +1 (*R*) or dies after being resighted (*R’*). The latter parameter is a nuisance parameter and is difficult to estimate. Including it in our model would add undue complexity given that most CARs occurred during peak spawning. Thus, we estimated our angler resighting parameter only during the summer survey occasions (*R_t_*).

As there was no expectation that survival or fishing mortality was the same across decades, and because no recaptures occurred between tagging periods, we fit separate models for each tagging period. We used beta distributions with flat priors for all four parameters (Table S1). We simulated data during model development to validate our ability to estimate true parameter values. For each model, we generated three Markov Chain Monte Carlo (MCMC) chains of 20,000 iterations, discarding the first 5,000 and saving every 5th iteration. We used a marginalized likelihood function to increase MCMC convergence speed (Yackulic et al. 2020), and we reported mean and Bayesian 95% credible intervals for each estimated and derived parameter (see sections 2.3.1 and 2.3.2 for derived quantities). We analyzed all CMR models in JAGS (Plummer 2003) with the R package `jagsUI` (Kellner 2021).

Given that most tagging in the 1960s and 1990s occurred during the summer, we opted for these models to be based on annual survey occasions, in which there was a single (Summer) tagging occasion per year (i.e., fish tagged outside of June-August were excluded from analysis). For the 2010s model, we modeled monthly survey occasions because tagging occurred year-round. As a result, we included a fixed effect of season (Summer, non-Summer) on survival, harvest, and CAR rates in the 2010s. We adjusted monthly harvest rates in the 2010s to annual harvest rates. Data filtering resulted in zero biologist recaptures and angler CARs in the 1990s and so the biologist recaptures (*p*) and CAR (*R*) parameters were fixed to zero in the 1990s model (Table S1).

#### 2.2.2 Tag Retention

To account for tag loss, we first separately modeled the probability of retaining a tag with data from a Kelp Bass (*Paralabrax clathratus*) double-tagging study that occurred off San Diego, CA, USA from 2012 to 2016 (see Bellquist 2015 for a detailed description of methods). We assumed tag retention rates to be similar between the two species and across tagging periods given that similar tagging methods were used by trained biologists in each of the three BSB tagging studies, and that Kelp Bass is a local congener of BSB with similar growth rates and overlapping habitat use (Lowe et al. 2003, Mason & Lowe 2010, Logan & Lowe 2018). We used a Bayesian hidden state framework in JAGS (Su & Yajima 2021, Plummer et al. 2022) to model tag retention over time at liberty, as a function of age of tag (Supplement S2).

Of the 673 Kelp Bass double-tagged in the tag retention experiment (Bellquist 2015), a total of 129 fish were recaptured within 3.7 years (31 with a single tag intact and 98 with both tags intact). The cumulative probability of (or proportion of fish) retaining at least one tag in the double tagging study was ∼ 86% in the first year and fell to 9% after seven years (Fig. S1a). Initial tag retention was estimated at ∼ 96% and the discrete annual rate of tag retention was estimated at ∼ 90%. The calculated probability (non-cumulative) of a fish retaining a tag, as a function of age of tag, decreased from ∼ 86% in the first year to ∼40% after five years and 34% after seven years (Fig. S1b).

The model estimated time-dependent probabilities of retaining a tag (*tr*, Fig. S1b) were incorporated into the CMR model framework for the 1960s and 1990s (see section 2.2.4). We defined the time-dependent tag retention priors with a beta distribution in which the shape parameters of each prior beta distribution in the CMR model were based on the mean and variance of the time-dependent tag retention posterior distributions derived from the tag retention model (Table S1). Given the short duration of the 2010s study (27 months) and that not all fish were tagged at the beginning of the study, we incorporated a prior for the discrete annual tag retention rate, exp(-*β*), termed *r* (Table S1).

#### 2.2.3 Growth Estimation

During the 1960s, 1990s, and through February 2013, the minimum size limit (MSL) was 305 mm (12 inches TL), corresponding to an average fishery recruitment age of five to six years (Love et al. 1996b); afterward, the MSL increased to 356 mm (14 inches TL, Jarvis et al. 2014), corresponding to an average fishery recruitment age of eight years (Walker et al. 2020b).

Our model accounts for potential harvest of sub-legal-size fish and CAR of legal-size fish. To do so, we incorporated growth in our model, such that at each time step (occasion), the size of each fish, if not supplied by the data, was estimated and the fish assigned a size class (sublegal or legal), whereby the probability of harvest (*κ*) and the probability of CAR (*R*) were estimated for both sublegal- and legal-size fish. To estimate BSB growth, we used the von Bertalanffy growth function (VBGF; Love et al. 1996b, Walker et al. 2020b). However, given that the parameters of the traditional VBGF are highly correlated (e.g., k, L_∞_; Ogle 2016), we instead used the Francis parameterization of the VBGF to estimate three growth parameters (L1, L2, and L3) in the CMR models. These parameters correspond to mean lengths at specific ages (Supplement S3). To generate priors for the Francis growth parameters in the CMR model, we separately modeled BSB growth using BSB age and growth data collected in southern California from 2011 to 2016 (Walker et al. 2020b) and the Francis parameterization of the VBGF in the R package FSA (Supplement S3, Ogle et al. 2022).

The 736 BSB collected in the age and growth study (Walker et al. 2020b) ranged in age from young-of-the-year (YOY) to 25 years, while total lengths ranged from 114 – 600 mm. The mean growth parameter estimates and 95% confidence intervals used to define priors in the CMR models were L1_3_ = 236 mm, CI: 229-242 mm (mean size at age 3 y); L2_9.5_ = 403 mm (mean size at age 9.5 y), CI: 400-406 mm; L3_16_ = 495 mm, CI: 487-502 mm (mean size at age 16 y, see Table S1 for priors). BSB males reach maturity between 2 and 5 years, and females between 2 and 5 years (Love et al. 1996b).

Given that fish growth in our model was informed by growth increments between recapture and tagging (or previous recapture events), which are independent of the size structure of the population, our model estimates of growth are robust to any fishing-influenced truncation in the length frequency distribution over time. Moreover, since we used the Francis parameterization of the VBGF to estimate growth, our estimates are directly comparable to estimates obtained by traditional age and growth studies using otolith increments (Francis 1988) and are more directly attributable to growth rate than if just sizes at age were sampled (Enberg et al. 2012).

#### 2.2.4. State-transition and observation matrices

Using a multi-state approach (Kéry & Schaub 2012), we defined the state transition matrix (S) to calculate the state transition probability for the three possible latent states in occasion *t+1* (columns), given the latent state in occasion *t* (rows): (1) alive with tag, (2) dead, and (3) unavailable for capture (dead by natural causes or lost tag),

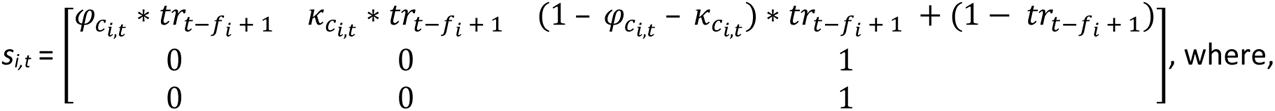

c*_i,t_* refers to the size class of individual *i* (legal, sublegal) at occasion t,

*tr_t – fi + 1_* is the tag retention probability at occasion *t + 1*, specific to the length of time the fish was at liberty (age of tag), where *f_i_* refers to the occasion of tagging. Note that for the 2010s model, instead of the time-varying *tr* parameter, we used a constant discrete annual probability of tag retention (*r*, Table S1) that was converted to a monthly rate (*r.mo.* = *r^1/12^*). In addition, the survival and harvest rate parameters were also indexed on season (spawning, nonspawning) at occasion *t* (e.g., 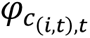).

We defined the observation matrix (O) to calculate the probability of observing each of five of the following possibilities in occasion *t+1* (columns), given the latent state in occasion t (rows): (1) recapture by a biologist and resighting by an angler, (2) recapture by a biologist, (3) resighting by an angler, (4) caught, kept, and reported by an angler, and (5), not seen or reported, where,

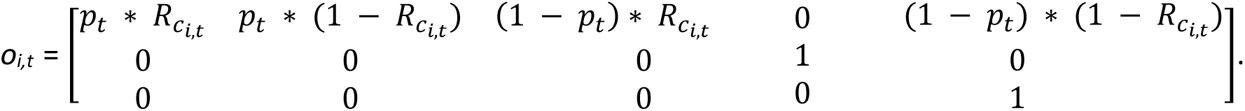

Note that for the 2010s model, the class and season indices were dealt with in the same manner as the state matrix above.

#### 2.2.5. Capture-history and Length Matrices

We constructed capture histories for each tagged fish and excluded fish tagged in the last survey occasion from the analysis. For a description of how we assigned recoveries to the correction survey occasion, see Supplement S1.

We constructed a length matrix consisting of lengths recorded at the occasion of release and those reported for angler resightings and angler recoveries. Given that our 1960s and 1990s CMR models estimated growth in annual increments, lengths of fish reported during the interval between survey occasions were assigned NAs. The length matrices for each period were supplied as data for the growth estimation portion of the CMR models.

### 2.3. Deriving Population Size

#### 2.3.1 Harvest Rates Chonditioned on Tag Reporting

The CMR model estimates of exploitation (i.e., harvest rate, *κ*) are dependent on tags of all resighted and recovered BSB being reported. When tag reporting is less than 100%, harvest estimates will be biased lower than the true harvest rate (Sackett & Catalano 2017). Given that tag reporting rates were unknown in this study and that reporting rates are known to vary widely across fisheries (Denson et al. 2002) we derived conditional size-specific harvest rates according to three hypothetical tag reporting probabilities of approximately 25%, 50%, and 75%. In this prior sensitivity analysis, for each tag reporting scenario, we assigned a corresponding beta distribution in the CMR model that we used *a posteriori*, in which we divided the posterior estimates of harvest rate by the prior distribution of the respective tag reporting. The assigned tag reporting priors included uncertainty of approximately ± 10% around the mean probability (Fig. S2).

#### 2.3.2. Population Size Conditioned on Tag Reporting

To calculate population estimates for each decade, we applied the mean conditional size-specific harvest rates to the annual size-specific harvest from each tagging period, where the annual legal size BSB harvest divided by the conditional harvest rate of legal size BSB equals the population of legal size BSB (see Supplement S4 on estimating size-specific harvest; the relative proportions of annual size-specific harvest are reported in Table S2). In the 1960s CMR, there was only a single year of size-specific catches for which to apply the size-specific conditional harvest rates, which yielded a single estimate of population size under each tag reporting scenario. In contrast, we were able to apply size-specific conditional harvest rates to the mean harvest across multiple years in the 1990s and 2010s to yield decadal estimates of population size under each tag reporting scenario.

To explore decadal trends in population size (and size-specific conditional harvest rates), we generated posterior distribution plots for each decade and tag reporting rate using the R packages tidybayes and ggdist (Kay 2022a,b).

### 2.4. Comparison to SST, Adult Densities, and Harvest

We obtained daily SST measurements (degrees Celsius) collected from 1954 to 2022 at the northern end of Santa Monica Bay, off Point Dume, CA, USA (Carter et al. 2022, Fig. 1). From these, we derived a mean summer (June-August) SST for each year. Of the coastal locations with available long-term, SST data in southern California (Carter et al. 2022), Pt. Dume is nearest to the location of the SCUBA transect data analyzed in this study (Fig. 1).

We obtained diver survey densities of adult (≥ 220 mm) and juvenile (< 150 mm, YOY) BSB from 1974 to 2022 in King Harbor, Redondo Beach, CA, USA (Fig. 1), collected by the Vantuna Research Group (VRG), Occidental College (unpublished data; see Stephens et al. 1986 for a detailed description of methods). Following an approximate pelagic larval duration of one lunar month (Allen & Block 2012), recruits settle into inshore nursery areas like bays and harbors (Love et al. 1996b). Once settled, recruits remain in these areas for about a year before moving deeper into the open coast; adults primarily associate with ecotone habitat during the non-spawning season and sand flats during the spawning season (Mason et al. 2010, McKinzie et al. 2014). Notably, the adult density data represent BSB from three years of age, which is approximately two-three years younger than most fishery recruits.

We plotted temporal trends in SST, adult densities, and harvest to compare their decadal means occurring during each tagging period and their overall patterns throughout the time series. Here, we were focused on trends and not absolute values (e.g., SST trends in Santa Monica Bay follow SST trends along the southern California coast). To identify potential lagged relationships between SST and adult density and between SST and harvest, we standardized each data set to a mean of zero and calculated cross-correlation coefficients from lags zero to ten years using the R package `funtimes` (Lyubchich et al. 2023). Positive correlations occurring at a lag of zero suggest influence of SST on the adult population, whereas positive correlations occurring at lags greater than three years (i.e., SST predicts future adult densities or harvest) suggest influence of SST on the early life history stages. The R package `funtimes` uses a bootstrap approach to account for potential autocorrelation among time series at each lag; of primary interest are correlations occurring near or outside the reported 95% confidence band.

### 2.5. Relationship Between Juvenile Recruitment and SST

To explore potential relationships between juvenile (YOY) recruitment and SST, we also considered the Ocean Niño Index (ONI), as El Niño was shown to have a positive effect on BSB larval abundances off Baja California (Avendaño-Ibarra et al. 2009). We obtained the ONI data as monthly index values (NOAA 2023a), which we averaged to obtain mean annual indices. We standardized both data sets to a mean of zero and tested for lagged correlations between SST and recruitment and between ONI and recruitment from lags zero to three years using the R package `funtimes` (Lyubchich et al. 2023). We further explored the influence of SST and ONI on juvenile recruitment with a generalized additive model (GAM) using the R package `mgcv` (Wood 2017). To account for possible confounding effects of spawning biomass and harvest impacts on juvenile recruitment, we incorporated adult densities and CPFV harvest in our model, but we excluded the period after the fishery collapse (after 2005), in which BSB CPFV harvest was consistently less than the historic minimum (88 thousand BSB in 1978). We specified a Tweedie observation error family (positive continuous density values that also contain zeros) and a log link, allowing the model to estimate the shape of the Tweedie distribution parameter. We specified all main effects as a penalized smooth function with a basis function (i.e., ‘wiggliness’) of three.

Based on the results of the cross-correlation analysis, we tested three candidate temperature models, 1) one with SST only (lag of zero), 2) one with SST and ONI (both with lag of zero), and 3) one with SST (lag of zero) and ONI (lag of one). In all models, adult densities and harvest were incorporated with a lag of zero. We performed model checks for convergence and basis function misspecification. We selected the most parsimonious model based on the lowest Akaiki information criterion (AIC) value and we report model fit as the percent deviance explained. We visually explored the conditional effects of important explanatory variables using the R package `visreg` (Breheny & Burchett 2019).

### 2.6. Historical accounts

Given limited species-specific harvest records and fishery-independent data prior to the mid-1970s, we gathered historical points of reference for BSB availability from the literature (see Supplement S5 for search terms). We compiled a table of BSB accounts spanning the mid-1800s through the late 1970s that referred to the relative contribution of BSB to commercial or recreational harvest, or that made any mention of BSB distribution, availability, or spawning in southern California. We then created a graphical timeline for contextualizing these accounts with respect to changes in BSB fishing regulations, the oceanographic climate, and trends in Rockbass CPFV harvest (the longest harvest time series that includes BSB). For the graphical timeline, we plotted monthly indices of the Pacific Decadal Oscillation (PDO, a measure of SST anomalies, NOAA 2023b) along with a 12-month running mean. We noted decadal-scale periods of predominately cool or warm temperature regimes (Minobe 1997, Mantua et al. 1997) associated with assemblage shifts in California’s fishes as described in Hubbs (1948), McCall (1996), and Overland et al. (2008). We also noted major El Niño events resulting in either seasonal warm water intrusions of subtropical and tropical fauna or decadal-scale northern range expansions of temperate/subtropical/tropical fauna in California (Hubbs 1948, Radovich 1961, Lea & Rosenblatt 2000, McClatchie 2014, Walker et al. 2020a).

### 2.7. Data and Code Availability

Data and code pertaining to the CMR model (simulation, growth, tag retention, CMR models by decade) are available online in a GitHub repository: https://github.com/ ETJarvisMason/bsb-CMR. We performed all analyses in R 4.0.3 (R-Core-Team 2020).

## 3. RESULTS

### 3.1. Decadal Trends in Demographic Rates

Mean annual BSB survival (*ϕ*) differed by size class (legal vs sublegal) and was higher for sublegal fish than legal-size fish, except in the 2010s (Fig. 2). By decade, mean annual survival was highest in the 1960s and lowest in the 1990s. Mean annual survival of legal-size BSB in the 2010s was higher than in the 1990s, but survival of sublegal fish in the 2010s was lower than in both the 1990s and the 1960s (Fig. 2). Overall, the estimated annual survival rates were substantially lower than would be expected based on the size distribution of fish tagged in the study (e.g., the size distribution of fish tagged included large BSB corresponding in age to 10+ years old, suggesting a much higher survival rate). Biologist recapture rates (*p*) of tagged BSB were low (≤ 1%) across all tagging periods. CAR rates could only be estimated for the 1960s and 2010s and were generally low (0-4%) compared to harvest rates (see section 3.2), but were nevertheless slightly conservative, as they assume 100% tag reporting.

**Figure 2.**
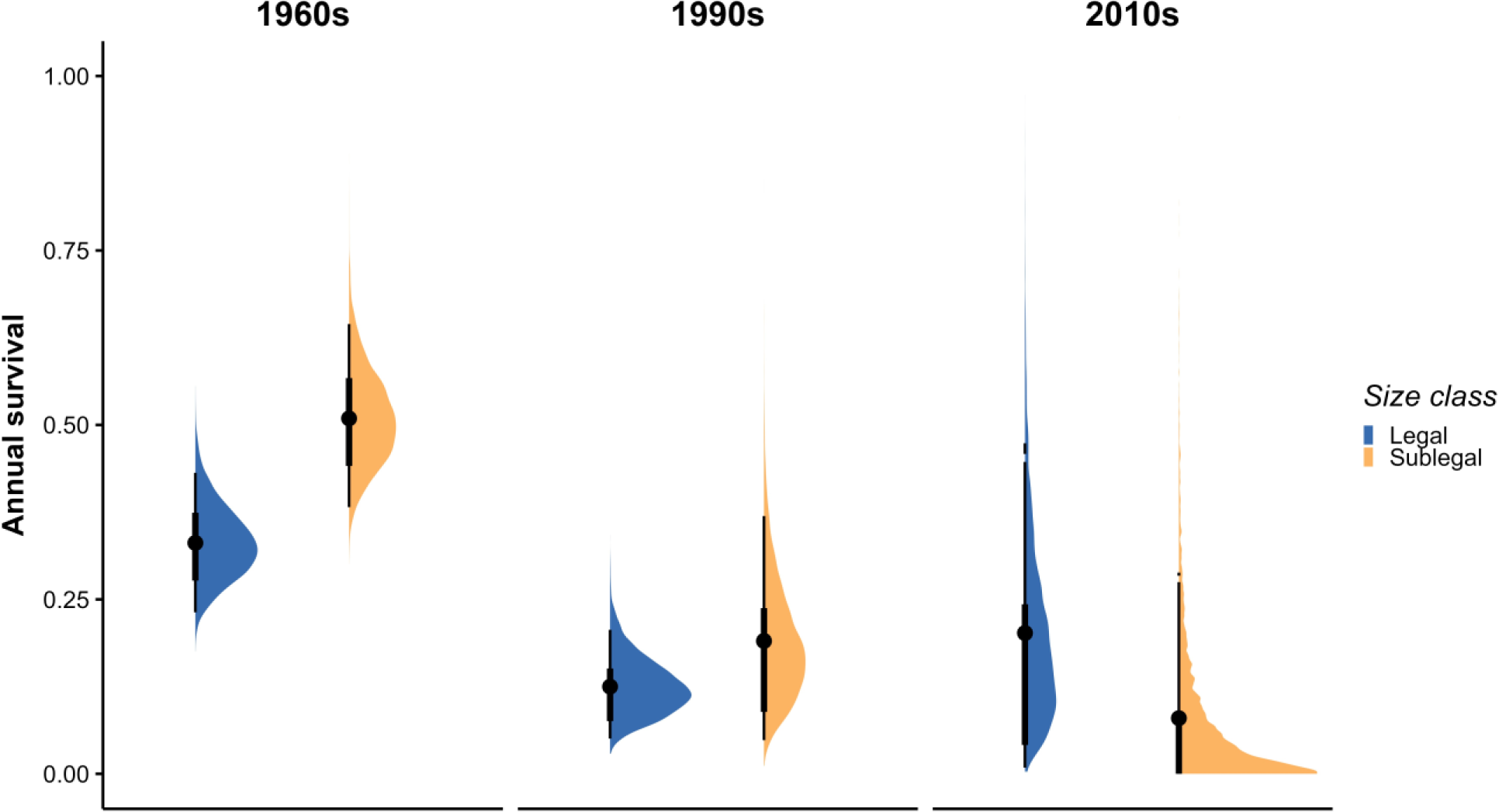
Bayesian capture-mark-reencounter model posterior distributions and mean annual survival and 66 and 95% HDI credible intervals (dots plus thick and thin lines) for legal- and sublegal-size Barred Sand Bass across tagging periods. Annual survival rate is the proportion surviving in a year.

There were too few recapture lengths from which to model growth in the 2010s; however, given that Walker et al. (2020b) collected BSB for age and growth during the same period (2011-2015), we included Francis VBGF growth parameter estimates from those data for comparison with the 1960s and 1990s CMR growth estimates. The CMR decadal growth estimates indicated BSB grew faster and reached a smaller size over time (Fig. 3).

**Figure 3.**
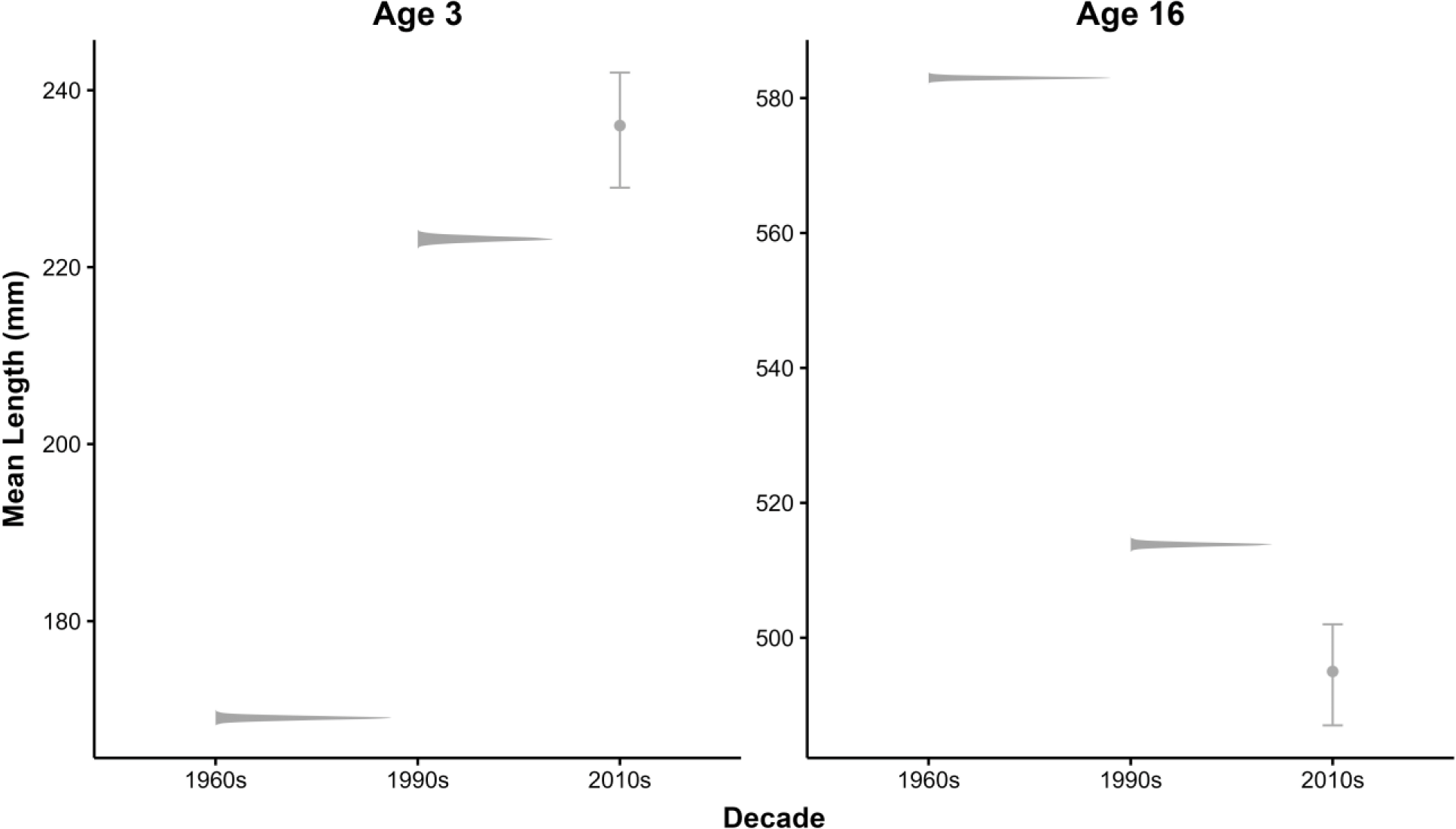
Parameter estimates of Barred Sand Bass mean lengths at ages 3 and 16 across tagging periods. The 1960s and 1990s growth parameter estimates represent capture-mark-reencounter model Bayesian posteriors for the Francis parameters, L1 and L3, of the von Bertalanffy growth function, while the 2010s estimates represent mean and 95% confidence intervals derived from Barred Sand Bass age and growth data collected from 2011 to 2016 and fit to the Francis parameterization of the von Bertalanffy growth function (there were too few recapture lengths in the 2010s data to accurately estimate growth in the 2010s capture-mark-reencounter model).

### 3.2. Decadal Trends in Harvest Rates and Population Size

Like the CAR rates, harvest rates estimated by our CMR model were also conservative because they are conditional on angler reporting rates. Without adjusting for tag reporting, we observed an overall decrease in legal-size harvest rates over time, with the 1960s harvest rate more than ∼2x and ∼5x higher than the 1990s and 2010s harvest rates, respectively (Fig. S2). Harvest rates of sublegal-size BSB were low across decades but increased slightly in the 2010s (Fig. S2). After adjusting for different tag reporting rates, with uncertainty, conditional harvest rates showed a similar pattern, regardless of tag reporting rate combination across decades. Harvest rates under a 25% reporting rate were highest but the most uncertain (Fig. 4a).

**Figure 4.**
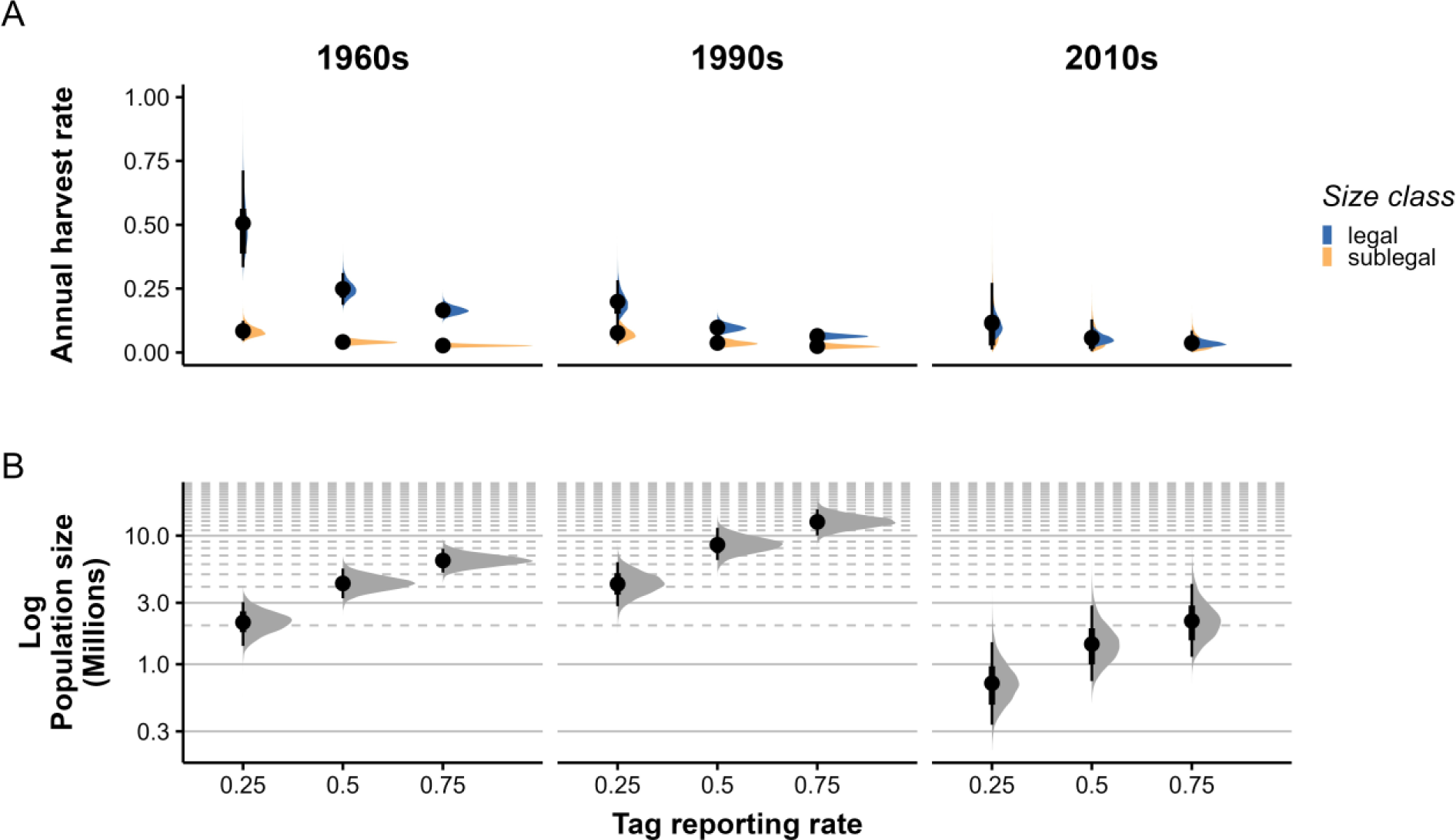
Bayesian capture-mark-reencounter model posterior distributions and 66 and 95% HDI credible intervals (dots plus thick and thin lines) of Barred Sand Bass a) size-specific annual harvest rates and b) log population size, conditioned on tag reporting scenario, with the following mean probabilities of reporting a tag, 25%, 50%, and 75%. Log population size in each decade was, in part, derived using the mean harvest of sublegal and legal-sized Barred Sand Bass across multiple years, except for the 1960s, in which only one year of harvest was available for estimating population sizes. Annual harvest rate is the proportion of fish dying due to fishing.

Ages obtained from 736 BSB collected in the age and growth study (southern California, 2011-2015; Walker et al. 2020b) ranged from YOY to 25 years, while total lengths ranged from 114 – 600 mm. The mean Francis VBGF parameter estimates and 95% confidence intervals estimated from these data and used to derive priors in the CMR models were L1_3_ = 236 mm, CI: 229-242 mm; L2_9.5_ = 403 mm, CI: 400-406 mm; L3_16_ = 495 mm, CI: 487-502 mm.

The CMR decadal growth estimates indicated BSB grew faster and reached a smaller size over time (Fig. S3). There were too few recapture lengths from which to model growth in the 2010s; however, given that (Walker et al. 2020b) collected BSB for age and growth during the same period (2011-2015), we included Francis VBGF growth parameter estimates from those data for comparison with the 1960s and 1990s CMR growth estimates.

The conditional estimates of mean BSB population size increased with increased tag reporting rates (Fig. 4b). Given the higher uncertainty in conditional harvest rates under the lowest probability tag reporting scenario (Fig. 4a), we focused our comparison of mean decadal population estimates under the 50% and 75% tag reporting scenarios (range: ∼ 40 – 85%).

Regardless of tag reporting combination across decades, the population increased by at least 50% to as much as double between the 1960s and 1990s (Fig. 4b). By the 2010s, the mean population size had declined nearly 10-fold to ∼ 1 to 3 million BSB. This was roughly 1/3 the size it had been in the 1960s, though there was greater uncertainty in the 2010s estimate (Fig. 4b). A posteriori, the maximum and minimum conditional mean population estimate across decades were ∼16 million BSB in the 1990s under a 75% tag reporting rate and ∼ 1.4 million BSB in the 2010s under a 50% tag reporting rate (Fig. 4b).

### 3.3. Comparison to SST, Adult Densities, and Harvest

Trends in our population estimates corresponded to trends in SST and fishery-independent and -dependent data during the same time periods (Fig. 5). Adult densities were not available prior to 1974, but the lower adult densities in the 1970s followed on from our relatively smaller population estimate in the 1960s (compared to the 1990s); adult densities were also lowest in the 2010s; Fig. 5b). Likewise, relative to the 1990s, BSB harvest by all fishing modes combined and by CPFVs alone was lower in the 1960s and lowest in the 2010s (Fig. 5c). Trends in BSB harvest were reflected in the large fluctuations in Rockbass (= Kelp Bass and Barred Sand Bass) CPFV harvest, corresponding to two windows of increased fishing opportunity for BSB (Fig. 5c). The first fluctuation in Rockbass harvest consisted of a substantial increase in the 1960s followed by a decline in the 1970s, and the second was an increase into the 1980s and 1990s followed by a precipitous decline in the 2000s. Overall, trends in adult densities and harvest appeared to lag SST trends; however after the fishery collapse, which occurred during the last tagging period in the mid-2010s, trends diverged, with SST and adult densities increasing, while BSB harvest remained low after 2015 (Fig. 5).

**Figure 5.**
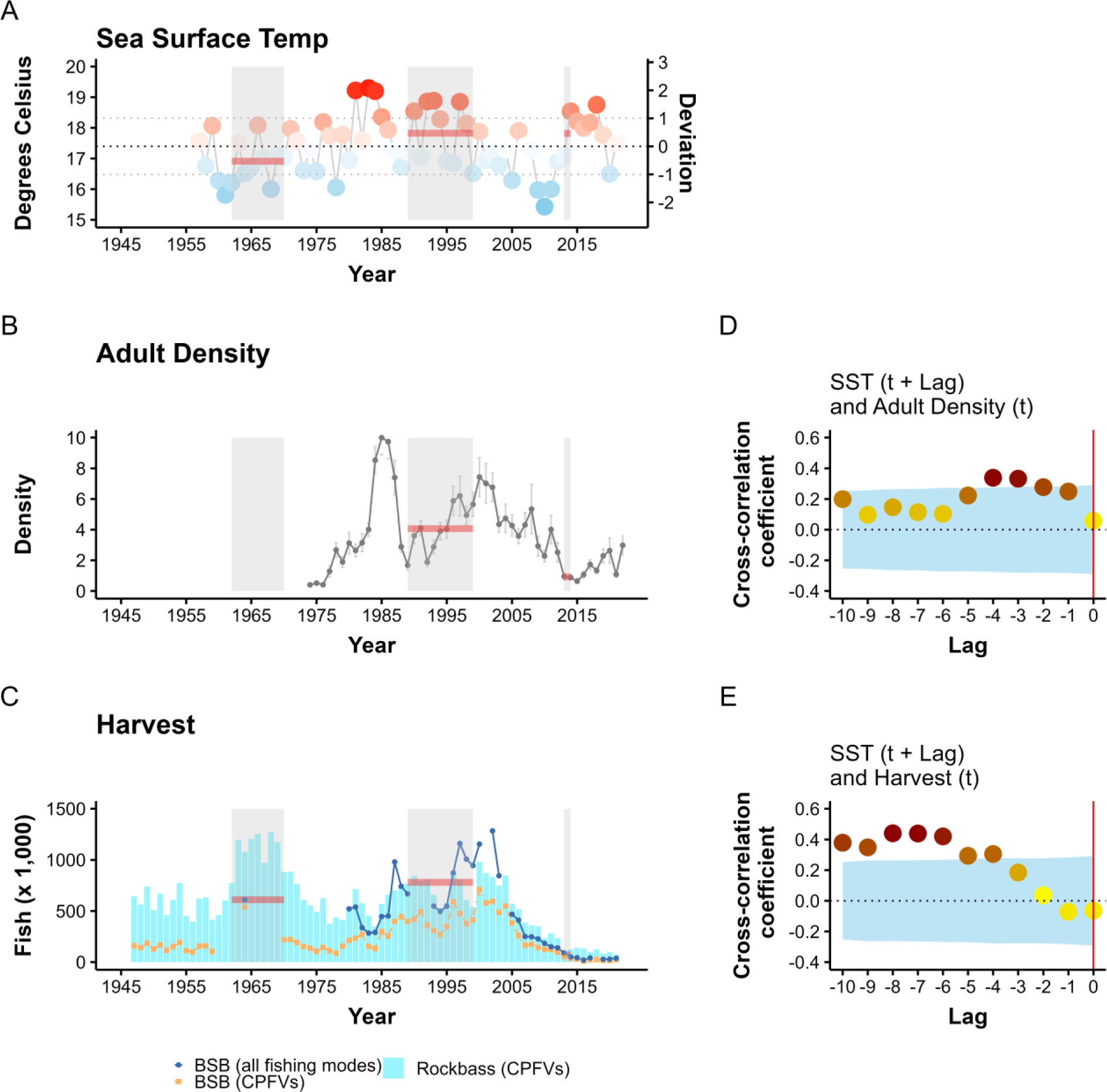
Temporal trends in a) average monthly summer sea surface temperatures (°C) at Pt. Dume, California, USA, 1956-2021, b) mean annual adult densities (fish/transect) of Barred Sand Bass as measured on diver surveys in King Harbor, California, USA, 1974-2022, c) total Barred Sand Bass harvest in southern California across all recreational fishing modes (dark blue solid line with closed circles; 1964 and 1980-2021), and cross-correlation coefficients for d) SST and lagged adult densities, and e) SST and lagged Commercial Passenger Fishing Vessel (CPFV) harvest of Barred Sand Bass (orange dashed dotted line with closed squares). CPFV harvest of Rockbass (= Kelp Bass and Barred Sand Bass, aqua bars) is included for reference. Horizontal lines depict means during each tagging period (shaded rectangular regions).

Cross-correlation analysis revealed adult densities lagged SST by three to four years, corresponding to the age at which BSB become mature (Fig. 5d), and harvest lagged SST by four to ten years, corresponding to ages of cohorts comprising the harvest data (Fig. 5e). The highest correlations between SST and harvest were strong and occurred at 6, 7, and 8 y, corresponding to the approximate age of fishery recruits. Though the adult densities represent data from a single location in southern California, the moderate to strong correlations identified by the cross-correlation analysis represent biologically meaningful lags between SST and adult densities, and between SST and harvest (Bight-wide data), suggesting the adult density data are representative of trends throughout the SCB.

### 3.4. Relationship Between Juvenile Recruitment and SST

During the 1990s tagging period, when the population size was estimated to be the highest, recruitment was generally below average while the mean SST was above average, and the ONI was mostly neutral except for the major El Niño event in 1997 (Fig. 6). Between 1974 and 2012, recruitment was sporadic, with only three of the 40 years showing strong peaks (one spanning the years 1977-79, one in 1984, and one in 1998); however, during the fishery collapse, from 2013 to 2021, BSB recruitment remained at elevated levels (Fig. 6c). Cross-correlation analysis revealed juvenile recruit density showed the highest correlation with SST at a lag of zero (Fig. 6d) and the highest correlation with the ONI at a lag of one (Fig. 6e). These correlations were only marginal and weak; however, the GAM revealed a strong positive relationship between recruitment and SST and recruitment and ONI, and the model fit was improved after incorporating a one-year lag on the ONI, with 36.3% of the deviance explained (Table 2). In contrast, the GAM revealed no effect of harvest and adult densities on juvenile BSB recruitment (Table 2). Given the potential that some YOY recruits may be closer in age to one than zero, we also tested the improved model with adult densities and harvest at a lag of one. This fourth model was the most parsimonious, revealing improved, but nevertheless marginal, effects of adult densities and harvest on BSB juvenile recruit densities compared to SST and the ONI (Table 2). When we considered the entire time series, in which adult densities and harvest were removed, the conditional effects plot showed an increasing, nonlinear positive effect of SST on BSB juvenile recruitment across increasing values of ONI, representative of La Niña, Neutral, and El Niño conditions (Fig. 6f).

**Figure 6.**
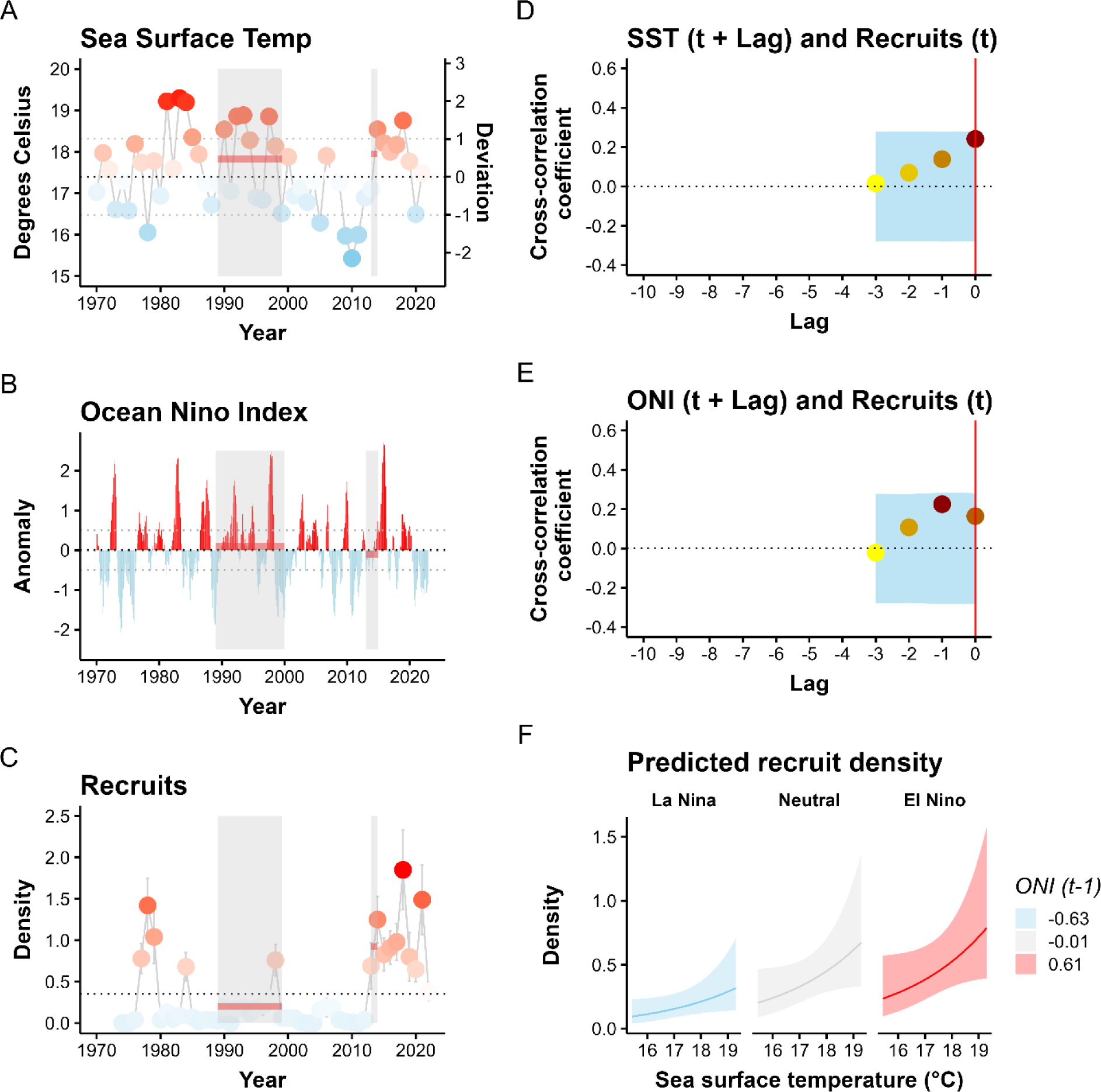
Temporal trends in a) average monthly summer sea surface temperatures (°C) at Pt. Dume, California, USA, 1970-2021, b) mean annual Ocean Niño Index anomalies c) mean annual densities of Barred Sand Bass juvenile (young-of-the-year) recruits as measured on diver surveys in King Harbor, California, USA, 1974-2022, c) cross-correlations of lagged SST and recruits, d) cross-correlations of lagged ONI and recruits, and d) conditional plot of the effect of SST on juvenile recruit density across different values of the ONI using all years of data. Horizontal red lines depict means during each tagging period (shaded gray regions).

**Table 2.**
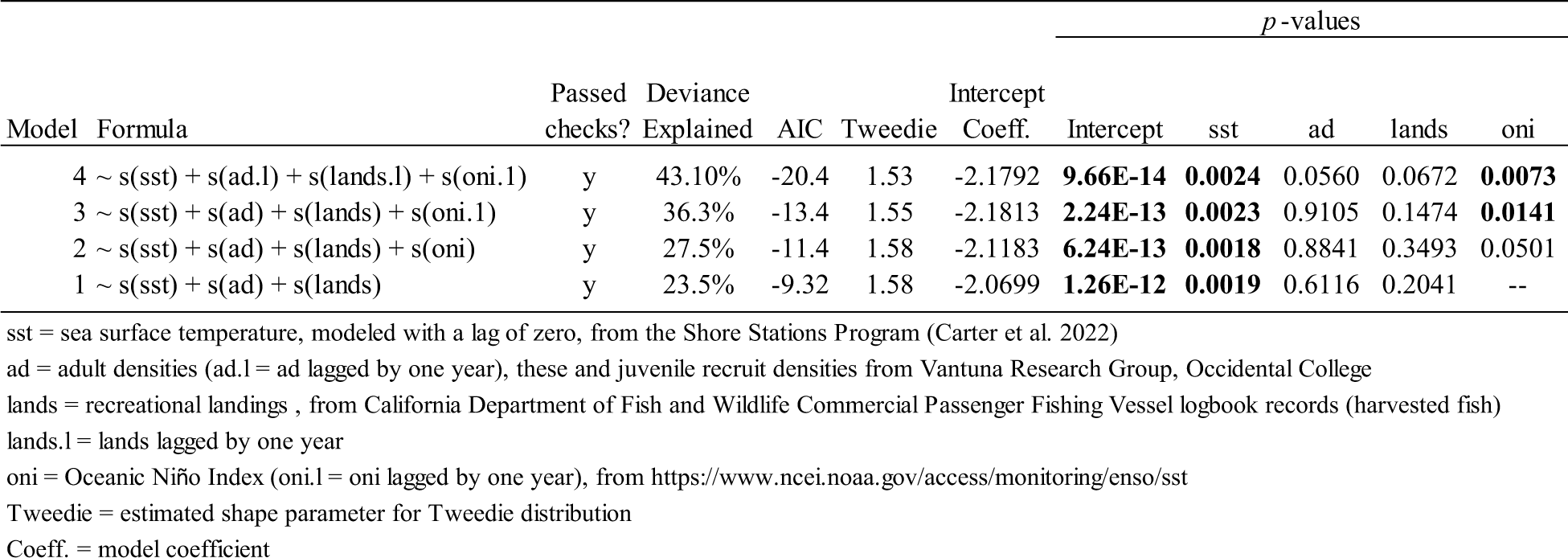
Results of the generalized additive model of juvenile (young-of-the-year) Barred Sand Bass recruit densities as a smoothed function of temperature (SST and ONI), adult densities, and harvest in southern California, USA, from 1974 to 2004.

### 3.5. Historical Accounts

Sources for BSB historical accounts included scientific journal publications (n = 16), a fishing guide, a publication on the status of California’s marine resources, and several government documents (n = 4) available online and by request, including CDFW administrative reports (n =3), and a CDFW monthly report (Fig. 7). When considered collectively, the historical accounts corresponded with the results of our quantitative analysis, in which periods of reportedly higher and lower BSB population abundance generally corresponded to decadal-scale fluctuations in ocean temperature (here, we used the PDO as a proxy for SST anomalies). Most notable were three periods, 1) the mid-19^th^ century, in which the southern California fish fauna was described as tropical and the distribution of BSB was documented as far north as Monterey in central California (Fig. 7a; Girard 1858, Hubbs 1948), 2) the cool regime of the 1950s and 1960s (Fig. 7f-h), in which BSB was referred to by CDFW field biologists as “scarce”, “a more southern species”, and comprising “a very small portion of the catch” relative to Kelp Bass (Young 1963, Young 1969, Feder et al. 1974), and 3) a short window in the 1960s (during the cool regime) when observations made by CDFW field biologists conducting diver and fishing surveys indicated a dramatic increase in the numbers of BSB in southern California (Fig. 7g). This observed increase in BSB availability was also reflected in the substantial increase in Rockbass harvest at the time (Fig. 7) and came on the heels of one of the most significant El Niño events documented in southern California (the 1957/58 El Niño; Fig. 7g). The higher Rockbass harvest was short-lived, and by the end of the cool regime, Rockbass harvest had dramatically declined and returned to being dominated by Kelp Bass (Fig. 5c, 7i; Wine 1978, 1979a,b). Love et al. (1996a) reported BSB CPFV CPUE at this time was 5-10x lower than it was a decade later, during the warm regime of the 1980s and 1990s (Fig. 7i;). For a more detailed narrative of BSB historical accounts, see Supplement S6.

**Figure 7.**
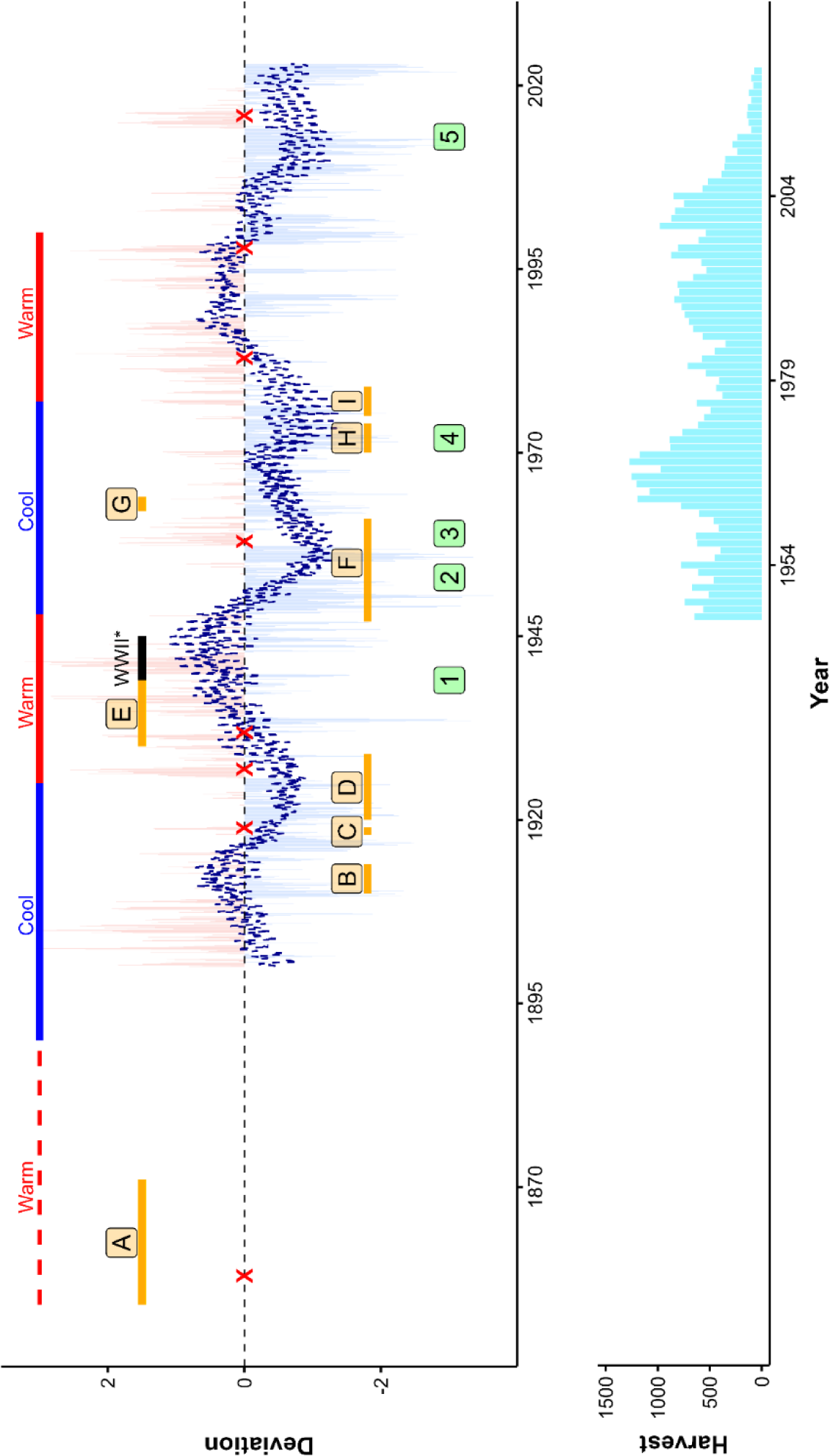
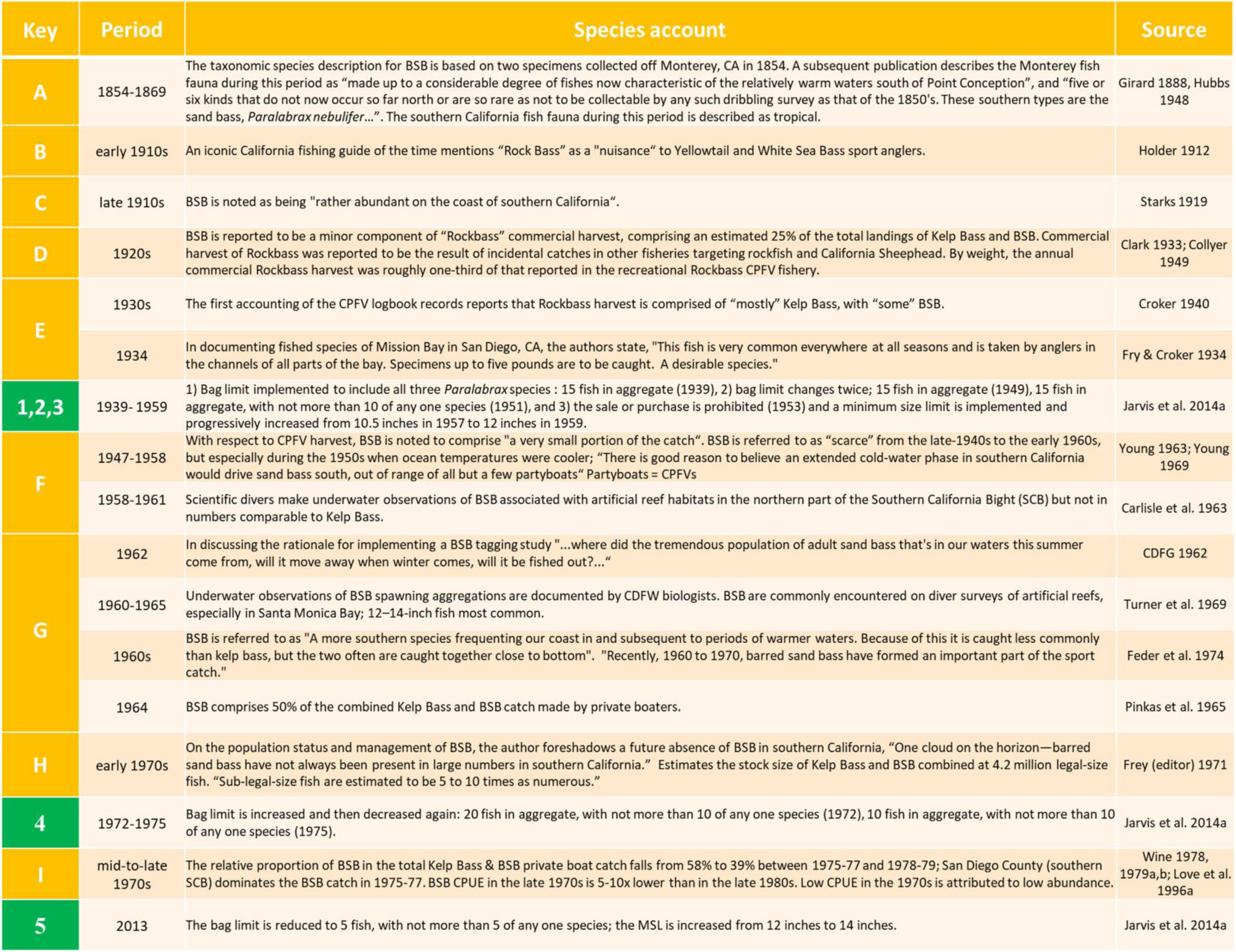
Graphical timeline (left) and trends in Rockbass CPFV harvest (thousands of fish, right) for contextualizing historical accounts of Barred Sand Bass harvest, distribution, and availability in California, USA, from the mid-nineteenth century to the 2020s (reference table, next page). Rockbass harvest includes Kelp Bass. The trend line represents a 12-month running average of the Pacific Decadal Oscillation, a measure of SST anomalies); periods designated as cool and warm are based on Minobe (1997) and Mantua et al. (1997). **X** = El Niño resulting in either seasonal warm water intrusions of subtropical and tropical fauna or decadal-scale northern range expansions of temperate/subtropical fauna in California. *No Commercial Passenger Fishing Vessel fishing permitted for five years during World War II. Reference table for the graphical timeline, representing historical accounts of Barred Sand Bass harvest, distribution, and availability (letters), and fishing regulation changes (numbers) in California, USA, from the mid-nineteenth century to the 2020s.

## 4. DISCUSSION

We have taken advantage of a multidecadal tag and recapture dataset to generate the first estimates of historical and contemporary BSB demographic rates and population size spanning different oceanographic regimes and harvest histories. Our estimates, combined with historical accounts in the literature and a variety of long-term data streams representing different life history stages, indicate the BSB fishery in southern California between 1962 and 2014 can be characterized by two windows of fishing opportunity that were largely driven by sporadic, warm-water recruitment events followed by efficient harvest on spawning aggregations. The last window resulted in a prolonged period of fishery collapse, in which we estimate the population had declined by approximately 1 order of magnitude. Despite this dramatically reduced population size and evidence of recruitment limitation in the 40 years prior, juvenile recruitment remained elevated in the decade since and was associated with novel, anomalously warm conditions. Thus, we can confirm that recruitment limitation was not a factor contributing to delayed recovery to-date. Based on these findings, we postulate that juvenile recruitment in southern California may not have been entirely locally sourced and that there is high potential for fishery recovery. Nevertheless, despite signs of incipient population recovery, the spawning aggregations remain absent, suggesting other factors have impacted fishery recovery.

### 4.1. Sporadic, warm-water recruitment pulses

At least since 1974, the BSB population in southern California has had extended periods of minimal juvenile (YOY) recruitment with intermittent peaks showing a strong relationship with SST, especially following El Niño events (Fig. 6c,d, Table 2). Moreover, this influence of SST on the early life history stage was detected in relationships between SST and future adult densities and harvest, implicating temperature as a major driver of future fishery recruitment. Our results are consistent with other studies; Miller & Erisman (2014) found that YOY BSB abundance from 1979 to 2010 was highly episodic, having a moderate positive relationship with SST and strong positive relationship with future CPUE in the fishery. In addition, Jarvis et al. (2014) found that *Paralabrax* spp. larval abundance and SST between 1996 and 2012 were positively correlated with future fishery recruitment strength.

Correspondence between early life history stage recruitment and fluctuations in future harvest/CPUE is characteristic of a population driven by recruitment limitation, in which varying recruitment levels are good predictors of subsequent population size (Armsworth 2002). This is noteworthy because periodic fluctuations in harvest/CPUE are generally atypical of aggregation-based fishery catch dynamics. Aggregate spawners typically show hyperstable catches, in which population declines are masked by stable catch rates as aggregation densities are maintained (Sadovy & Domeier 2005, Erisman et al. 2011). For example, among overexploited fisheries, “plateau-shaped” harvest trajectories are common in hyperstable fisheries (“i.e., a sudden fall after a relatively long and stable persistence of high level catches”, Mullon et al. 2005); however, BSB showed a more “erratic” harvest trajectory (“i.e., a fall after several ups and downs”, Mullon et al. 2005). We focused here on harvest rather than CPUE because there was longer availability of harvest trends and because in this fishery, harvest trends closely correspond with CPUE trends through time (Jarvis et al. 2014, CDFW 2020). Although effort shifts can contribute to interannual fluctuations in recreational harvest (Dotson and Charter 2003, Blincow and Semmens 2022), our analysis nevertheless revealed a strong relationship between SST and harvest at biologically meaningful lags.

Juvenile recruitment has remained well above average since 2013, despite the low population size estimated for the 2010s when the fishery collapsed. Between 2012 and 2020, southern California experienced several marine heatwaves (MHW, 2014-15 [“the Blob”], 2019, 2020, 2021), including a strong El Niño (2015-16). The effects of this dramatic alteration of the Southern California Bight ecosystem were profound (Leising 2015, Cavole et al. 2016, Walker et al. 2020a) and, in some cases, atypical of expectation based on previously established environment-species relationships (McClatchie et al. 2018, Thompson et al. 2019, 2022). This anomalous warm water is likely to have had a positive effect on any locally sourced BSB larvae and may have also resulted in externally sourced BSB larvae from Baja California. Although MHWs lack the strong northward horizontal transport characteristic of El Niño (Amaya et al. 2020), they can result in an “abrupt diminishing of upwelling” off Baja California (Jiménez-Quiroz et al. 2019), thereby eliminating any barrier to northward larval transport that is typically present during the summer months. An example of this potential is a record of an adult Goldspotted Sand Bass (*P. auroguttatus*, a congener rare north of Baja California Sur), that was first documented in southern California off Santa Barbara in 2018 (Love et al. 2019), four years following “the Blob”.

Given the sporadic nature of BSB juvenile recruitment and positive correlation with ONI, it is possible that the southern California population is dependent on El Niño-driven larval transport from Baja California (Lilly et al. 2022). El Niño is known to facilitate poleward advection of larvae into southern California (McClatchie et al. 2018, Cimino et al. 2021, Lilly et al. 2022). Indeed, genetic connectivity exists between BSB populations in the two regions (Paterson et al. 2015) and recruitment dependence on Baja California fish populations has been suggested for other fishes in southern California (Smith & Moser 1988, Allen & Franklin 1992, Ben-Aderet et al. 2020). In contrast to BSB, the Kelp Bass population in southern California, which has more reliably persisted, was found to be locally sourced (Selkoe et al. 2007). The southern California BSB population is at the northern extent of its core population range, and recruitment is typically more variable for marine populations at their geographic margins (Myers 1991, Neill et al. 1994, Levin et al. 1997).

Relative to the BSB spawning season in southern California, the spawning season off Baja California is more protracted (May through Feb), with a summer and fall peak in larval abundance and higher abundance during El Niño events (Avendaño-Ibarra et al. 2009). Thus, following an El Niño year, a portion of YOY recruits in southern California may represent northward advected Baja California larvae from the previous summer or fall, which would correspond to the one-year lag we observed between the ONI and juvenile densities. Nevertheless, the degree to which BSB larval recruitment is seeded from Baja California warrants further research.

### 4.2. Impacts to Aggregation Dynamics?

Following sustained recent juvenile recruitment, adult densities increased to more than double the levels prior to 2015; however, BSB harvest has remained exceptionally low. Under the current MSL, BSB are expected to recruit to the fishery at an average age of eight years; thus, the earliest indication of fishery recovery should have been evident in 2020. One explanation for the lack of fishery recruitment despite high juvenile recruitment and higher adult densities, could simply be that it is still too early to detect in the harvest data, as boat access to ocean fishing was halted during the COVID-19 pandemic and some commercial sportfishing vessel operations were also temporarily closed. We also cannot rule out the possibility of a potential residual Allee effect associated with the population decline and fishery collapse in the mid-2010s. For example, in healthy transient aggregate spawner populations, the permanence of spawning aggregation locations is maintained by social transmission over many generations (e.g., older adults know where to go from experience and younger adults learn by following older adults, Warner 1988). If harvest removes enough of the older adults or densities are low enough, social transmission is interrupted. This may result in many smaller localized aggregations or the establishment of new aggregation sites at locations unknown to anglers (Warner 1988, Waterhouse et al. 2020). An acoustic telemetry study off San Diego, CA between 2012 and 2016 showed evidence of adult BSB spawning season migrations to a previously undocumented aggregation site, however, the larger traditional spawning grounds never manifested aggregations (Bellquist 2015).

### 4.3. Trends in Demographic Rates

Between the 1960s and 1990s, technological advances in locating aggregations afforded greater precision in targeting spawning sites (Allen & Hovey 2001), and so we expected a higher exploitation rate in the 1990s. However, conditional harvest rates were generally higher in the 1960s and lower in the 1990s, regardless of tag reporting rate. This could be in part due to a higher number of licensed anglers in the 1960s than in the 1990s (∼3x more, Bellquist 2015). In addition, the mean annual harvest of sublegal- and legal-sized fish in both decades was similar (Table S2), and our CMR model results indicate the BSB population size in the 1990s was bigger relative to the 1960s. Thus, even though targeting spawning aggregations may have become easier by the 1990s, the sizable increase in BSB population size would have resulted in a smaller fraction of BSB being removed due to fishing, despite increased harvest efficiency.

Harvest rates were most uncertain under the 25% tag reporting scenario. Although tag reporting rates were unknown for each tagging period, we assume they ranged from moderate (∼50%) to high (∼75%). Of the three conditional harvest rates for the 1990s, the harvest rate under a mean 50% probability of reporting a tag was most similar to the exploitation rates reported for the 1990s using catch curves (∼11%, Jarvis et al. 2014). We presume that tag reporting in the 1960s was at least as high or higher due to enhanced outreach and cooperation with the fishing community at that time (Collyer & Young 1953, Young 1963). It is possible that cooperation to report tags was not as high in the 2010s, given the increased take restrictions in 2013.

Based on the size structure of the tagged populations in each of the three tagging decades, we are certain our model estimates of survival are biased low, which could be due to 1) decreased tag reporting over time (due to e.g., faded ink, excessive biogenic growth on tags; Waterhouse & Hoenig (2012) or 2) invalid assumptions regarding fidelity of tagged BSB to the southern California tagging area (e.g., we assumed no permanent emigration; Barker 1997). The latter is much less likely since BSB home ranges are small and the average migration distance to spawning grounds in southern California is ∼ 15 km (Jarvis et al. 2010, Mason & Lowe 2010). Despite this bias, the trend in our survival estimates (highest in the 1960s, lowest in the 1990s) suggests that conditions in the 1990s were less favorable to adult BSB survival even though exploitation was lower. The slight increase in survival of legal-size BSB in the 2010s coincided with the implementation of tighter fishing regulations in 2013, while survival of sublegal-size BSB in the 2010s was lowest of the three tagging periods. This lower survival rate may have also contributed to a lack of fishery recovery, though the mechanism is unclear.

Our model estimates of BSB growth in the 1960s are the first published historical estimates prior to the 1990s (Love et al. 1996b). We detected directional changes in the mean-size-at age through time, in which the magnitude of change was greater between the 1960s and 1990s than between the 1990s and 2010s); BSB grew slightly faster by age 3 and grew slower by age 16 (Fig. 3). Fish growth rates can show high phenotypic plasticity resulting from the environment (e.g., temperature, food availability), density-dependent processes, and fishing. However, when larger, older fish are predominantly harvested, changes to growth and maturity can result from fishing-induced evolution (Enberg et al. 2012). Just prior to the fishery collapse in the mid-2000s, BSB catches switched from being dominated by young adult fishery recruits to older, larger fish (Jarvis et al. 2014), but BSB size and age at maturity have not been re-evaluated since the 1990s (Love et al. 1996b).

### 4.4. Shifting Baselines

Historical ecology is a valuable tool that can increase our understanding of the factors influencing fluctuations in populations and consequently improve our ability to evaluate a population’s potential for decline and recovery (Scarborough et al. 2022). In this study, when considered collectively, the historical accounts of BSB that we gathered also served to validate the results of our quantitative analysis; that is, periods of reportedly higher and lower BSB population abundance were associated with decadal-scale fluctuations in ocean temperature.

One notable finding was that observations of increased BSB availability in the 1960s were reflected in the substantial increase in Rockbass (= Kelp Bass and Barred Sand Bass) CPFV harvest during that period (Fig. 5c). Prior to this study, the increase in Rockbass harvest in the 1960s could not be attributed to Kelp Bass or BSB based on CPFV logbook records alone due to inconsistencies in species-specific reporting prior to 1975. However, additional catch survey data we gathered from that period indicate the contribution of BSB to Rockbass harvest doubled relative to previous estimates (Pinkas et al.1968, Fig. 5c, 7g). The dramatic increase in BSB availability occurred five years after the strong 1957-58 El Niño, a period corresponding to the approximate age when BSB recruit into the fishery (at the time, age five to six years) and one that further supports our findings relating sporadic juvenile recruitment pulses to warm water events. It was also critical in discovering this fishery experienced more than one boom and bust period.

Given the high fishing mortality in the 1960s and apparent recruitment limitation in the southern California BSB population, it is not surprising that harvest quickly returned to low levels by the mid-1970s. The decrease in availability was correctly foreshadowed by resource managers (Frey 1971) and yet, they did not express concern, as they had come to expect lower BSB abundance during cooler conditions. Thus, a “healthy” BSB population is likely to look different to different people, depending on the lifetime of perspective (Bellquist et al. 2017). We found that the exceptional increased availability of BSB during the warm regime of the 1980s and 1990s was not the norm for much of the last century. Shifting baselines and institutional amnesia can result in diminished expectations of what the size of a healthy aggregate spawner population should be, inadvertently resulting in less conservative, less effective, management measures (Fulton 2023).

### 4.5. Management Implications

Given the relationship we identified between recruitment and warm water pulses, we might anticipate that the predicted increase in MHWs (Oliver 2019) and secular ocean warming would benefit the BSB population in the future with perhaps more consistent strong recruitment and a shift in the center of the BSB geographic distribution northward into southern California (Pinsky et al. 2020). Although recent trends in juvenile recruitment suggest fishery recovery is imminent, current regulations may not be adequate to prevent the quick collapse of a new emerging cohort and thus, management preparedness is prudent. Nassau Grouper (*Epinephelus striatus*), which also exhibits sporadic pulse recruitment (Stock et al. 2021, 2023), showed evidence of recovery 15 years following conservation measures that included seasonal closures (Waterhouse et al. 2020). A seasonal closure for BSB could potentially serve to enhance local recruitment, especially during favorable oceanographic conditions, and combined with the existing bag limit and MSL, limit the potential for subsequent collapse and delayed recovery.

Our population estimates suggest that the prolonged fishery collapse following the second window of BSB fishing opportunity in the last century resulted from an approximate 10-fold decline in the population (Fig. 4). The first window of fishing opportunity in the 1960s did not result in a similar prolonged collapse. One difference between the two periods is that temperatures during the catch declines that led to the second collapse remained cooler longer and there was no major El Niño event for nearly a decade (Fig. 5,7). There is also no evidence to suggest that illegal fishing (private boaters or shore anglers taking more than the bag limit) or underreporting of catch (on CPFV logbooks) has increased over time. Additionally, the reduced population size during the second fishery collapse may have resulted in adult densities reaching an Allee-effect threshold (i.e., levels that could prolong collapse or delay/impede recovery, Hutchings 2015, Sadovy de Mitcheson 2016). If true, then a potential Allee-effect threshold for BSB, based on our CMR decadal population estimates of adult and subadult BSB (legal- and sublegal-sized BSB), could be somewhere around 3 million fish. Moreover, we found that juvenile (YOY) recruitment was more influenced by warm water pulses than adult densities or harvest, suggesting recruitment is less tied to spawning stock biomass than might otherwise be assumed. This breakdown, or lack of a spawner-recruit relationship is not uncommon (Lowerre-Barbieri, Heyman 2019), and should be explored further if a formal stock assessment is considered for BSB.

Our results demonstrate that even with long-standing harvest limits in place (e.g., minimum size limit, bag limit), aggregation-based fishing in recruitment-limited populations may force the fishery to exist in perpetual boom-and-bust. Such a model of fishing opportunity is unwise for recreational fisheries that are known to have considerable social and economic benefits (Griffiths et al. 2017, Lovell et al. 2020) and are intended to be sustainable for future generations. Thus, if the goal is to maintain reliable recreational fishing opportunities for a recruitment-limited transient aggregate spawner, limiting high CPUE fishing opportunities, i.e., limiting fishing during the spawning season, may be the best insurance against sudden, prolonged collapse and would balance the protection of spawning aggregations with long-term fishery sustainability (Erisman et al. 2020). Our results demonstrate the importance of historical context and long-term monitoring in resolving the role of sporadic recruitment and aggregation-based fishing in driving the population dynamics of an iconic aggregate spawner.

## ACKNOWLEDGEMENTS

This research was supported with a QUEST grant awarded by NOAA Fisheries (NOAA) to B. Semmens. We thank C. Valle and H. Gliniak (CDFW) for the historical BSB tag and recapture data and age and growth data, J. Laake (retired, NOAA) and O. Gimenez (Centre d’Ecologie Fonctionnelle et Evolutive) for their statistical expertise, and A. Thompson (NOAA) for constructive comments on an earlier draft of the manuscript. We are grateful to the late J. Stephens (founder of VRG in 1974) for his dedication to VRG’s long-term SCUBA monitoring program.

## LITERATURE CITED

Aguilar-Perera A (2006) Disappearance of a Nassau grouper spawning aggregation off the southern Mexican Caribbean coast. Mar Ecol Prog Ser 327:289–296.

Allee W (1931) Animal Aggregations, a Study in General Sociology. University of Chicago Press, Chicago, USA.

Allee W (1938) The Social Life of Animals, First Edition. WW Norton Inc., New York, USA.

Allen L.G, Block HE (2012) Planktonic Larval Duration, Settlement, and Growth Rates of the Young-of-the-Year of Two Sand Basses (*Paralabrax nebulifer* and *P. maculatofasciatus*: fam. Serranidae) from Southern California. Bull South Calif Acad Sci 111:15–21. 10.3160/0038-3872-111.1.15

Allen LG, Franklin MP (1992) Abundance, distribution, and settlement of young-of-the-year white seabass *Atractoscion nobilis* in the Southern California Bight, 1988-89. Fish Bull 90:633–641.

Allen LG, Hovey TE (2001) Barred sand bass. In: California’s living marine resources: a status report. Leet WS, Dewees CM, Klingbeil R, Larson EJ (eds) Department of Fish and Game, Sacramento, USA, p 224–225

Armsworth PR (2002) Recruitment limitation, population regulation, and larval connectivity in reef fish metapopulations. Ecology 83:1092–1104.

Avendaño-Ibarra R, Hernández-Rivas ME, de Silva-Dávila R (2009) Reproductive Strategies of Sea Basses based on Larval Abundance in Magdalena Bay, Mexico, 1982–1986. North Am J Fish Manag 29:205–215.

Barker RJ (1997) Joint modeling of live-recapture, tag-resight, and tag-recovery data. Biometrics 53:666–677.

Bellquist LF, Semmens BX, Stohs S, Siddall A (2017) Impacts of recently implemented recreational fisheries regulations on the Commercial Passenger Fishing Vessel fishery for *Paralabrax* sp. In California. Mar Policy 86:134–143.

Bellquist LF (2015) A historical perspective of California recreational fisheries using a new database of ‘trophy’ fish records (1966-2013), combined with fisheries analyses of three species in the genus Paralabrax. PhD dissertation, University of California, San Diego, CA

Ben-Aderet N, Johnston EM, Cravey R, Sandin SA (2020) Revisiting the life history of yellowtail jack (*Seriola dorsalis*) in the Southern California Bight: New evidence for ontogenetic habitat shifts and regional differences in a changing environment. Fish Bull 118:157–170.

Blincow KM, Semmens BX (2022) The effect of sea surface temperature on the structure and connectivity of species landings interaction networks in a multispecies recreational fishery. 1119:1109–1119.

Bolden SK (2000) Long-distance movement of a Nassau grouper (*Epinephelus striatus*) to a spawning aggregation in the central Bajamas. Fish Bull 96:642–644.

Breheny P, Burchett W (2019) Visualization of Regression Models Using visreg. R J 9:56–71.

CDFG [California Department of Fish and Game] (1962) Report for the month of August, 1962. Dept. of Fish and Game Records, Regional Reports, F3498:582–614.

Carlisle J, Turner C, Ebert E (1963) Artificial Habitat in the Marine Environment. Fish Bull 124.

Carter ML, Flick, Reinhard E, Terrill E, Beckhaus, Elena C. Martin K, Fey CL, Walker, Patricia W. Largier, John L. McGowan JA (2022) Shore Stations Program Data Archive: Current and historical coastal ocean temperature and salinity measurements from California stations. UC San Diego Library Digital Collections. 10.6075/J00001XZ

Chollett I, Priest M, Fulton S, Heyman WD (2020) Should we protect extirpated fish spawning aggregation sites? Biol Conserv 241:108395.

Cimino MA, Jacox MG, Bograd SJ, Brodie S, Carroll G, Hazen EL, Lavaniegos BE, Morales MM, Satterthwaite E, Rykaczewski RR (2021) Anomalous poleward advection facilitates episodic range expansions of pelagic red crabs in the eastern North Pacific. Limnol Oceanogr 66:3176– 3189.

Clark FN (1933) Rock bass (*Paralabrax*) in the California commercial fishery. Calif Fish Game 19:25–35.

Collyer R (1949) The Commercial Fish Catch of California for the Year 1947: With an Historical Review 1916-1947. Fish Bull:268.

Collyer R & Young P H (1953) Progress Report on a Study of the Kelp Bass, *Paralabrax clathratus*. Calif Fish Game 39:191–208.

Croker RS (1940) Three Years of Fisheries Statistics on Marine Sport Fishing in California. Trans Am Fish Soc 69:111–118.

Denson MR, Jenkins WE, Woodward AG, Smith TIJ (2002) Tag-reporting levels for red drum (*Sciaenops ocellatus*) caught by anglers in South Carolina and Georgia estuaries. Fish Bull 100:35–41.

Domeier ML, Colin PL (1997) Tropical reef fish spawning aggregations: defined and reviewed. Bull Mar Sci 60:698–726.

Dotson RC, Charter RL (2003) Trends in the southern California sport fishery. CalCOFI Reports 44:94–106.

Enberg K, Jørgensen C, Dunlop ES, Varpe Ø, Boukal DS, Baulier L, Eliassen S, Heino M (2012) Fishing-induced evolution of growth concepts mechanisms and the empirical evidence. Mar Ecol 33:1–25.

Erisman BE, Allen LG, Claisse JT, Pondella DJ, Miller EF, Murray JH, Walters C (2011) The illusion of plenty: hyperstability masks collapses in two recreational fisheries that target fish spawning aggregations. Can J Fish Aquat Sci 68:1705–1716.

Erisman BE, Grüss A, Mascareñas-Osorio I, Lícon-González H, Johnson AF, López-Sagástegui C (2020) Balancing conservation and utilization in spawning aggregation fisheries : a trade-off analysis of an overexploited marine fish. ICES J Mar Sci 77:148–161.

Feder HM, Turner CH, Limbaugh C (1974) Observations On Fishes Associated With Kelp Beds in Southern California. Calif Dep Fish Game, Fish Bull 160.

Francis RICC (1988) Are growth parameters estimated from tagging and age-length data comparable? Can J Fish Aquat Sci 45:936–942.

Frey HW (ed) (1971) Kelp and sand bass In: California’s Living Marine Resources and Their Utilization. California Department of Fish and Game, Sacramento, USA, p 93–94

Fulton S (2023) Institutional amnesia pushes fish spawning aggregations towards extirpation. People Nat 5:489–495.

Fry D, Croker RS (1934) A preliminary survey of Mission Bay State Park. Calif Fish Game 20:1– 13.

Girard C (1858) Fishes. In: *General report upon the zoology of the several Pacific railroad routes*. Explorations and surveys for a railroad route from the Mississippi River to the Pacific Ocean. 10(4):1–400.

Griffiths SP, Bryant J, Raymond HF, Newcombe PA (2017) Quantifying subjective human dimensions of recreational fishing: does good health come to those who bait? Fish Fish 18:171–184.

Heemstra PC (1995) Serranidae. Meros, serranos, guasetas, enjambres, baquetas, indios, loros, gallinas, cabrillas, garropas. In: Guia FAO para Identification de Especies para lo Fines de la Pesca. Pacifico Centro-Oriental. 3 Vols. Fischer W, Krupp F, Schneider W, Sommer C, Carpenter KE, Niem V (eds) FAO, Rome, p 1565–1613

Heyman WD, Grüss A, Biggs C, Kobara S, Farmer N, Karnauskas M, Lowerre-Barbieri S, Erisman B (2019) Cooperative monitoring, assessment, and management of fish spawning aggregations and associated fisheries in the U.S. Gulf of Mexico. Marine Policy, 109: 103689.

Hilborn R, Hively DJ, Jensen OP, Branch TA (2014) The dynamics of fish populations at low abundance and prospects for rebuilding and recovery. 71:2141–2151.

Holder CF (1912) The fishes of the Pacific coast, a handbook for sportsmen and tourists. Dodge Publishing Company, New York, USA.

Hsieh CH, Reiss C, Watson W, Allen MJ, Hunter JR, Lea RN, Rosenblatt RH, Smith PE, Sugihara G (2005) A comparison of long-term trends and variability in populations of larvae of exploited and unexploited fishes in the Southern California region: A community approach. Prog Oceanogr 67:160–185.

Hubbs C (1948) Changes in the fish fauna of western north america correlated with changes in ocean temperature. J Mar Res VII: 459–482.

Hutchings J A (2015) Thresholds for impaired species recovery. In Proceedings of the Royal Society B: Biological Sciences (Vol. 282, Issue 1809). Royal Society of London. 10.1098/rspb.2015.0654

Jarvis ET, Gliniak HL, Valle CF (2014a) Effects of fishing and the environment on the long-term sustainability of the recreational saltwater bass fishery in southern California. Calif Fish Game 100:234–259.

Jarvis ET, Linardich C, Valle CF (2010) Spawning-Related Movements of Barred Sand Bass, *Paralabrax nebulifer*, in Southern California: Interpretations from Two Decades of Historical Tag and Recapture Data. Bull South Calif Acad Sci 109:123–143.

Kay M (2022a) ggdist: Visualizations of Distributions and Uncertainty. R package version 3.2.0

Kay M (2022b) Tidybayes: Tidy Data and Geoms for Bayesian Models. R package version 3.0.2

Kellner K (2021) JagsUI: a wrapper around “rjags” to streamline “JAGS” analyses. R package version 1.4.2

Kéry M, Schaub M (2012) Chapter 9 – Estimation of Survival and Movement from Capture– Recapture Data Using Multistate Models. In: Bayesian Population Analysis using WinBUGS. Kéry M, Schaub M (eds) Academic Press, p 263–313

Kuparinen A, Keith DM, Hutchings JA (2014) Increased environmentally driven recruitment variability decreases resilience to fishing and increases uncertainty of recovery. ICES J Mar Sci 71:1507–1514.

Lea RN, Rosenblatt RH (2000) Observations on fishes associated with the 1997-98 El Niño off California. CalCOFI Reports 41:117–129.

Levin PS, Chiasson W, Green JM (1997) Geographic differences in recruitment and population structure of a temperate reef fish. Mar Ecol Prog Ser 161:23–35.

Liermann M, Hilborn R (2001) Depensation: evidence, models and implications. Fish Fish 2:33– 58.

Lilly LE, Cornuelle BD, Ohman MD (2022) Using a Lagrangian particle tracking model to evaluate impacts of El Niño-related advection on euphausiids in the southern California Current System. Deep Res Part I Oceanogr Res Pap 187.

Logan RK, Lowe CG (2018) Residency and inter-reef connectivity of three gamefishes between natural reefs and a large mitigation artificial reef. Mar Ecol Prog Ser 593:111–126.

Di Lorenzo E, Schneider N, Cobb KM, Franks PJS, Chhak K, Miller AJ, McWilliams JC, Bograd SJ, Arango H, Curchitser E, Powell TM, Rivière P (2008) North Pacific Gyre Oscillation links ocean climate and ecosystem change. Geophys Res Lett 35:2–7.

Love MS, Brooks A, Ally JRR (1996a) An analysis of commercial passenger fishing vessel fisheries for kelp bass and barred sand bass in the Southern California Bight. Calif Fish Game 82:105–121.

Love MS, Brooks A, Busatto D, Stephens J, Gregory PA (1996b) Aspects of the life histories of the kelp bass, *Paralabrax clathratus*, and barred sand bass, *P. nebulifer*, from the southern California Bight. Fish Bull 94:472–481.

Love MS, McCrea M, Johnston D, Butterfield A (2019) First Authenticated Record of the Goldspotted Sand Bass, *Paralabrax auroguttatus* from California Waters. Bull South Calif Acad Sci.

Love MS, Passerelli JK (2020) Miller and Lea’s Guide to the Coastal Marine Fishes of California, 2nd ed. University of California Agriculture and Natural Resources.

Lovell SJ, Hilger J, Rollins E, Olsen NA, Steinback S (2020) The Economic Contribution of Marine Angler Expenditures on Fishing Trips in the United States, 2017. NOAA Technical Memorandum NMFS-F/SPO-201.

Lowe CG, Topping DT, Cartamil DP, Papastamatiou YP (2003) Movement patterns, home range, and habitat utilization of adult kelp bass. Mar Ecol Prog Ser 256:205–216.

Lowerre-Barbieri S, Crabtree L, Switzer T, Burnsed SW, Guenther C (2015) Assessing reproductive resilience: An example with South Atlantic red snapper *Lutjanus campechanus*. Mar Ecol Prog Ser 526:125–141.

Lowerre-Barbieri S, DeCelles G, Pepin P, Catalán IA, Muhling B, Erisman BE, Cadrin SX, Ospina-Alvarez A, Stachura MM, Tringali MD, Burnsed SW, Paris CB (2017) Reproductive resilience: a paradigm shift in understanding spawner-recruit systems in exploited marine fish. Fish Fish 18:285–312.

Lyubchich V, Gel Y, Vishwakarma S (2023) Package Funtimes: Functions for Time Series Analysis version 9.1

Mantua NJ, Hare SR, Zhang Y, Wallace JM, Francis RC (1997) A Pacific Interdecadal Climate Oscillation with Impacts on Salmon Production. Bull Am Meteorol Soc 78:1069–1079.

Mason TJ, Lowe CG (2010) Home range, habitat use, and site fidelity of barred sand bass within a southern California marine protected area. Fish Res 106:93–101.

McClatchie S (2014) Regional Fisheries Oceanography of the California Current System. Springer Netherlands, Dordrecht.

McClatchie S, Gao J, Drenkard EJ, Thompson AR, Watson W, Ciannelli L, Bograd SJ, Thorson JT (2018) Interannual and Secular Variability of Larvae of Mesopelagic and Forage Fishes in the Southern California Current System. J Geophys Res Ocean 123:6277–6295.

McKinzie MK, Jarvis ET, Lowe CG (2014) Fine-scale horizontal and vertical movement of barred sand bass, *Paralabrax nebulifer*, during spawning and non-spawning seasons. Fish Res 150:66–75.

Miller EF, Erisman B (2014) Long-term trends of southern California’s kelp and barred sand bass populations: A fishery-independent assessment. CalCOFI Reports 55:1–9.

Minobe S (1997) A 50–70 year climatic oscillation over the North Pacific and North America. Geophys Res Lett 24:683–686.

MacCall AD (1996) Patterns of low-frequency variability in fish populations of the California Current. CalCOFI Reports 37:100–110.

Moser HG, Charter RL, Smith PE, Ambrose D a, Watson W, Charter SR, Sandknop EM (2001) Distributional atlas of fish larvae and eggs in the Southern California Bight region: 1951-1998. CalCOFI Atlas 34:1951–1998.

Mullon C, Fréon P, Cury P (2005) The dynamics of collapse in world fisheries. Fish Fish 6:111– 120.

Myers RA (1991) Recruitment variability and range of three fish species. NAFO Sci Counc Stud 16:21–24.

Neill WH, Miller JM, Van Der Veer HW, Winemiller KO (1994) Ecophysiology of marine fish. Netherlands J Sea Res 32:135–152.

NOAA (2023a) El Niño/Southern Oscillation (ENSO). National Centers for Environmental Information. https://www.ncei.noaa.gov/access/monitoring/enso/sst (accessed 9 February 2023)

NOAA (2023b) Pacific Decadal Oscillation. National Centers for Environmental Information https://www.ncei.noaa.gov/access/monitoring/pdo/ (accessed 9 Jan 2023)

Ogle DH (2016) Introductory Fisheries Analyses with R. CRC Press.

Ogle DH, Doll J, Wheeler P (2022) Package ‘FSA’ version 0.9.3

Oliver ECJ (2019) Mean warming not variability drives marine heatwave trends. Clim Dyn 53:1653–1659.

Overland J, Rodionov S, Minobe S, Bond N (2008) North Pacific regime shifts: Definitions, issues and recent transitions. Prog Oceanogr 77:92–102.

Paterson CN, Chabot CL, Robertson JM, Erisman B, Cota-Nieto JJ, Allen LG (2015) The genetic diversity and population structure of barred sand bass, Paralabrax nebulifer: A historically important fisheries species off southern and Baja California. CalCOFI Reports 56:1–13.

Perälä T, Hutchings JA, Kuparinen A (2022) Allee effects and the Allee-effect zone in northwest Atlantic cod. Biol Lett 18:3–8.

Pine WE, Pollock KH, Hightower JE, Kwak TJ, Rice JA (2003) A Review of Tagging Methods for Estimating Fish Population Size and Components of Mortality. Fish Res 28:10–23.

Pinkas MS, Oliphant CW, Haugen L (1968) Southern California marine sport-fishing survey: private boats, 1964; shoreline, 1965-66. Calif Dep Fish Game, Fish Bull 143

Pinsky ML, Selden RL, Kitchel ZJ (2020) Climate-Driven Shifts in Marine Species Ranges: Scaling from Organisms to Communities. Ann Rev Mar Sci 12:153–179.

Plummer M (2003) JAGS: A program for analysis of Bayesian models using Gibbs sampling. Proc 3rd Int Work Distrib Stat Comput Vienna, Austria.

Plummer M, Stukalov A, Denwood M (2022) Package ‘rjags’ version 4-13.

Pondella DJ, Allen LG (2008) The decline and recovery of four predatory fishes from the Southern California Bight. Mar Biol 154:307–313.

R-Core-Team (2020) R: A language and environment for statistical computing.

Radovich J (1961) Relationships of Some Marine Organisms of the Northeast Pacific to Water Temperatures Particularly During 1957 Through 1959. Calif Dep Fish Game, Fish Bull 112

Riecke T V., Gibson D, Leach AG, Lindberg MS, Schaub M, Sedinger JS (2021) Bayesian mark– recapture–resight–recovery models: increasing user flexibility in the BUGS language. Ecosphere 12:1–10.

Roedel PM (1953) Common Ocean Fishes of the California Coast. Calif Dep Fish Game, Fish Bull:91

Sackett DK, Catalano M (2017) Spatial heterogeneity, variable rewards, tag loss, and tagging mortality affect the performance of mark–recapture designs to estimate exploitation: An example using red snapper in the northern Gulf of Mexico. North Am J Fish Manag 37:558– 573.

Sadovy Y, Domeier M (2005) Are aggregation-fisheries sustainable? Reef fish fisheries as a case study. Coral Reefs 24:254–262.

Sadovy Y, Eklund a. M (1999) Synopsis of biological data on the Nassau grouper, *Epinephelus striatus* (Bloch, 1792), and the jewfish, *E. itajara* (Lichenstein, 1822). Fao Fish Synopsis:68.

Sadovy de Mitcheson Y (2016) Mainstreaming Fish Spawning Aggregations into Fishery Management Calls for a Precautionary Approach. Bioscience 66:295–306.

Scarborough C, Welch S, Wilson J, Gleason MG, Saccomanno VR, Halpern BS (2022) The historical ecology of coastal California. Ocean Coast Manag 230:1–16.

Selkoe KA, Vogel A, Gaines SD (2007) Effects of ephemeral circulation on recruitment and connectivity of nearshore fish populations spanning Southern and Baja California. Mar Ecol Prog Ser 351:209–220.

Semmens B, Bush P, Heppell S, Johnson B, McCoy C, Pattengill-Semmens C, Waylen L (2008) Charting a Course for Nassau Grouper Recovery in the Caribbean: What We’ve Learned and What We Still Need to Know. In: Proceedings of the 60^th^ Gulf and Caribbean Fisheries Institute, November 5 – 9, 2007, Punta Cana, Dominican Republic. Gulf and Caribbean Fisheries Institute, Marathon, FL, USA, p 607–609.

Smith PE, Moser HG (1988) CalCOFI time series: an overview of fishes. CalCOFI Rep 29:66–78.

Stephens JS, Jr., Jordan GA, Morris PA, Singer MM, McGowen GE (1986) Can we relate larval fish abundance to recruitment or population stability? A preliminary analysis of recruitment to a temperate rocky reef. CalCOFI Rep 27:65–83.

Stephens JS, Morris PA, Pondella DJ, Koonce TA, Jordan GA (1994) Overview of the Dynamics of an Urban Artificial. Bull Mar Sci 55:1224–1239.

Stephens PA, Sutherland WJ, Freckleton RP (1999) What Is the Allee Effect? Oikos 87:185–190.

Starks EC (1919) The basses and bass-like fishes of California. Families Serranidae, Haemulidae, Kyphosidae. Calif Fish Game 5:56–68.

Stock BC, Heppell SA, Waterhouse L, Dove IC, Pattengill-Semmens C V., McCoy CM, Bush PG, Ebanks-Petrie G, Semmens BX (2021) Pulse recruitment and recovery of Cayman Islands Nassau Grouper (*Epinephelus striatus*) spawning aggregations revealed by in situ length-frequency data. ICES J Mar Sci 78:277–292.

Su Y-S, Yajima M (2021) Package ‘R2jags’ version 0.5-7

Szuwalski CS, Vert-Pre KA, Punt AE, Branch TA, Hilborn, R (2015) Examining common assumptions about recruitment: A meta-analysis of recruitment dynamics for worldwide marine fisheries. Fish and Fisheries, 16(4), 633–648. 10.1111/faf.12083

Teesdale GN, Wolfe BW, Lowe CG (2015) Patterns of home ranging, site fidelity, and seasonal spawning migration of barred sand bass caught within the Palos Verdes Shelf Superfund Site. Mar Ecol Prog Ser 539:255–269.

Turner CH, Ebert EE, Given RR (1969) Man-Made Reef Ecology. Calif Dep Fish Game, Fish Bull 146

Vert-Pre KA, Amoroso RO, Jensen OP, Hilborn R (2013) Frequency and intensity of productivity regime shifts in marine fish stocks. Proc Natl Acad Sci U S A 110:1779–1784.

Walker HJ, Hastings PA, Hyde JR, Lea RN, Snodgrass OE, Bellquist LF (2020a) Unusual occurrences of fishes in the Southern California Current System during the warm water period of 2014–2018. Estuar Coast Shelf Sci.

Walker KM, Pentilla KM, Jarvis-Mason ET, Valle CF (2020b) Validated age and growth of Barred Sand Bass within the Southern California Bight. Calif Fish Wildl J 106:205–220.

Warner RR (1990) Male versus female influences on mating-site determination in a coral reef fish. Anim Behav 39:540–548.

Warner RR (1988) Traditionality of mating-site preferences in a coral reef fish. Nature 335:719– 721.

Waterhouse L, Heppell SA, Pattengill-Semmens CV., McCoy C, Bush P, Johnson BC, Semmens BX (2020) Recovery of critically endangered Nassau grouper (Epinephelus striatus) in the Cayman Islands following targeted conservation actions. Proc Natl Acad Sci U S A 117:1587–1595.

Waterhouse L, Hoenig JM (2012) Tagging models for estimating survival rates when tag visibility changes over time: Partial-year tabulation of recaptures. North Am J Fish Manag 32:147–158.

Wine V (1978) Southern California Independent Sport Fishing Survey Annual Report No. 2. Mar Resour Adm Rep No 78-2:84.

Wine V (1979a) Southern California Independent Sport Fishing Survey Annual Report No. 3. Mar Resour Adm Rep No 79-3:105.

Wine V (1979b) Southern California Marine Sport Fishing: Private-Boat Catch and Effort, 1975-1976. Mar Resour Adm Rep No 79-11:64.

Wood S (2017). Generalized Additive Models: An Introduction with R, 2nd edition. Chapman and Hall/CRC.

