## Supplemental Material for "Recruitment limitation increases susceptibility to fishing-induced collapse in a spawning aggregation fishery"

### S1. *Data Formatting and Processing*

#### S1.1. *Recaptures*

In the 1960s and 1990s, the disposition of reencountered fish (i.e., kept or released) was not always provided. Unless otherwise reported, we assumed all sublegal fish recaptured by anglers were released and all legal-size fish were retained (but see section 2.2.3 in the Methods). During the 1960s, 1990s, and through February 2013, the minimum size limit (MSL) was 305 mm (12 inches TL), corresponding to a fishery recruitment age of five to six years (Love et al. 1996b); afterward the MSL increased to 356 mm (14 inches TL, Jarvis et al. 2014), corresponding to fishery recruitment at approximately eight years (Walker et al. 2020b). Records with unknown tagging lengths were removed from the analysis.

We assumed a reencountered fish was recaptured by a biologist if the recapture occurred on the same date and at the same location as a survey occasion *and* the recapture length was not missing. We estimated missing dates based on time at liberty (in years) calculated from the difference in age between tag and recapture events using published von Bertalanffy age and growth parameters from the 1990s (Love et al. 1996a). If a tagging date was missing and there was also a missing recapture length, the tagging date was deduced based on sampling dates at the tag location and the sequence of tag identification numbers at that location.

We identified outliers in the reported lengths of recaptured Barred Sand Bass (BSB) by first calculating growth increments of recaptured fish and standardizing them by time at liberty in years (mm $yr^{-1}$). We then examined the distributions of growth increments over fish lengths in 50 mm TL bins from 250 – 600 mm. We flagged negative growth and increments greater than 150 cm in a year as outliers (n = 84) and replaced the corresponding reported recapture lengths with NA.

S1.2. *Assignment of Recovery Occasions*

In the 1960s and 1990s data, if a fish was tagged the year prior and caught and kept the following year, but before the next survey occasion (e.g., Jan-May), then the fish did not survive the interval from $t$ to $t$+1, and we recorded the recovery observation as occurring in that same year ($t$+1). However, if a fish was tagged the year prior and caught and kept the following year during or after the following survey occasion (i.e., Jun-Dec), then the fish survived the interval from $t$ to $t$+1 and we recorded the recovery observation as occurring in the subsequent year, $t$+2. Thus, we pushed recovery occasions out by one occasion unless the fish was recovered before June. In the 2010s data, we pushed all recovery occasions out by one occasion since the non-survey interval was sufficiently short (less than one month).

S2. *Tag Retention Model*

We used a Bayesian hidden state framework in JAGS (Su & Yajima 2021, Plummer et al. 2022) and Kelp Bass double-tagging data (Bellquist 2015) to model BSB tag retention over time, as a function of age of tag:

$$Q_i = \alpha * \exp(-(\beta * t^{\gamma})), \text{ where,}$$

$Q_i$ is the fish-specific probability of retaining a tag after recapture interval $t$,

$\alpha$ is the probability of retaining a tag immediately after release,

$\beta$ is the continuous rate of long-term (chronic) tag loss (note that exp($-\beta$) is the discrete rate of retention in a single time step),

$t$ is the time at liberty, and

$\gamma$ is an exponent on time to account for age of tag.

The model accounts for the probability of a fish retaining both tags or just one tag, where,

$p_1 = (1 - Q_i) * Q_i + Q_i * (1 - Q_i)$ is the probability of retaining the first tag and losing the second tag or losing the first tag and retaining the second tag, and

$p_2 = 1 - p_1$ is the probability of retaining both tags.

The likelihood of the data was then drawn from a binomial distribution of one trial with probability equal to $p_2$. We first derived posterior estimates of the cumulative tag retention ($Q_t$) over time from one to ten years at liberty (the maximum number of survey occasions). This represented the mean proportion of fish in the double-tagging study still retaining at least one tag at each time step. We then solved for the mean time-dependent probabilities of retaining a tag with the following equation:

$$tr_t = 1- (Q_{t-1} - Q_t)/Q_{t-1}, \text{ where,}$$

$Q_{t-1}$ is the proportion of fish still retaining at least one tag at time $t-1$, and

$Q_t$ is the proportion of fish still retaining at least one tag at time $t$.

S3. *Growth Model*

We used the Francis parameterization of the Von Bertalanffy Growth Function (VFGF) in the R package FSA (Ogle et al. 2022) and Barred Sand Bass age and growth data from Walker et al. (2020) to estimate BSB growth parameters,

$$E[L|t] = L1 + (L3 - L1)\,\frac{1 - r^{2\frac{t-t_1}{t_3-t_1}}}{1 - r^2}$$, where,

$E[L|t]$ represents the estimated length at age,

$L1$, $L2$, and $L3$ are the mean lengths at ages $t_1$, $t_2$, and $t_3$,

$t_1$ and $t_3$ are not estimated but are assigned to correspond to “young” and “old” ages, respectively,

$t_2 = t_1+t_3 / 2$, and

$r = L3 - L2 / L2 - L1$.

We used the length parameter estimates generated from the Francis parameterization of the VBGF to define the priors in our CMR models (Table S1). For the 1960s and 1990s, we used length parameter estimates based on $t_1$ = 3 years and $t_3$ = 16 years. In the 2010s model, we modeled monthly growth, and thus, length priors were based on the age of fish in months: $t_1$ = 24 months and $t_3$ = 192 months (Table S1). We chose a smaller age at $t_1$ for the 2010s model because the minimum size tagged was smaller than in the 1960s and 1990s.

S4. *Size-specific Estimates of Annual Harvest*

Harvest includes fish caught and kept by Commercial Passenger Fishing Vessels (CPFVs), private boaters, and shore anglers. We obtained harvest in numbers from California

Department of Fish and Wildlife (CDFW) CPFV logbooks between 1947 and 2021. Harvest for BSB prior to 1975 was likely underreported because although catches of "sand bass" could be recorded in catch logs, Kelp Bass was the only *Paralabrax* species that was pre-printed on the logs for entering catch, and captains were not required to distinguish the bass species in their records (Croker 1940, Young 1969). Historically, CDFW biologists estimated BSB comprised a small portion of the bass catch through the 1950s (~25%; Clark 1933, Roedel 1953, Young 1969), but this increased to ~50% by at least the mid-1960s (Pinkas et al. 1968) and returned to 25% by at least the mid-1970s. We applied these percentages to the overall numbers of harvested bass (historically referred to as "Rockbass") reported in the logbooks to calculate estimated annual BSB CPFV catches prior to 1975.

Estimates of private boat and shore-based harvest from the 1960s were only available from 1964-65 (July-June, private boat) and 1965 (January-December, shore-based; Pinkas et al. 1968; Table S2). Thus, for the 1960s we have a single estimate of BSB harvest, in which the CPFV estimate from 1964 and the estimated private boat and shoreline catch were combined. For the other two decades, given that the CPFV logbook data is the longest running record of recreational bass harvest, we chose to account for other methods of BSB take by adjusting the annual BSB CPFV harvest. To do so, we added numbers equivalent to the proportion of private and shore-based BSB harvest in each year, according to the relative proportion of harvest by fishing modes available from southern California recreational survey estimates, which are based on angler-intercept and telephone surveys (1980-2003: Marine Fisheries Statistical Survey [MRFSS], 2005-2017: California Recreational Fisheries Survey [CRFS]; Table S2). Thus, the adjusted BSB harvest, which represents the total estimated harvest of BSB, was then comparable across years.

For each year of available harvest, we estimated the proportion of sublegal and legal-size fish harvested. For 1964, we multiplied BSB harvest by the proportion of sublegal and legal-size harvested BSB measured in recreational angler-intercept surveys by CDFW biologists in the year 1975 (Wine 1978), as this was the earliest year for which length data in the recreational harvest was available (Table S2). For the 1990s and 2010s, the annual harvest was split into sublegal and legal size by multiplying the total annual harvest by the relative annual proportions of both size classes obtained from recreational angler surveys (Table S2). We applied these size-specific estimates of annual harvest (sublegal and legal) to CMR model estimates of exploitation to derive size-specific estimates of population size during each tagging period (see *Harvest Rates* and *Population Size*).

S5. *Search Terms for Historical Literature Review*

We conducted our historical literature review on the Web of Science search engine, as well as with Google Scholar. Search terms included "Barred Sand Bass," "Sand Bass," "Sandbass," "*Paralabrax nebulifer*," "rockbass," and "rock bass."

S6. *Detailed Narrative of Historical BSB Accounts*

Between the 1850s and 1970s there were two extended warm periods in southern California, USA: the first warm period was from 1854 to 1870 and the second was from about 1925 to 1947; however, relative to the first warm period and the warm period in the 1980s and 1990s, the second warm period was only moderately warm (Fig. 7; Hubbs 1948, McClatchie

2014). BSB was first taxonomically described in 1854 during the first warm period, when its distribution was documented as far north as Monterey, CA (Fig. 7a; Girard 1858, Hubbs 1948). In the early 20th century, accounts of BSB in the literature shifted from no mention to being noted as a minor species in California's early commercial fishery (Fig. 7b,c,d). During the moderately warm period (1925-1947), CDFW biologists estimated the commercial Rockbass harvest consisted of 25% BSB and 75% Kelp Bass. Most commercial Rockbass were incidentally taken when fishing for other species (i.e., rockfish, California Sheephead); by weight, recreational Rockbass harvest was three times higher than commercial Rockbass harvest.

Between 1920 and 1939, CPFV fishing became more affordable and by 1936, a catch logbook was required to be submitted (Fig. 7e). Shortly after, in 1939, a bag limit for the three saltwater basses of 15 fish in aggregate was implemented. The early description of the combined CPFV bass harvest was "mostly" Kelp Bass, with "some" BSB (Croker 1940). During the warmest part of the moderately warm period in the mid-1940s, there was a five-year reprieve from CPFV fishing due to World War II, and thus, no catch records exist (Young 1969). Following the war, the oceanographic climate shifted to a cold regime, during which BSB were reportedly "scarce" and comprised a "very small portion of the catch" (Fig. 7f; Young 1963, Young 1969). In the 1950s, a series of sportfishing regulations were implemented for the basses due to concerns over the Kelp Bass resource and declining catches (Fig. 7; Jarvis et al. 2014).

In 1962, CDFW field biologists noted "tremendous" numbers of BSB in southern California waters and initiated the BSB tagging study from which our model results are drawn (Fig. 7g; CDFG 1962). This apparent dramatic increase in BSB availability was also referenced in Young (1969), Frey (1971), and Feder et al. (1974), and was reflected in the substantial increase in Rockbass harvest during the 1960s (Fig. 5c). The 1962 increase in availability came on the heels of one of the most significant El Niño events documented in southern California (the 1957/58 El Niño; Fig. 7g). However, compared to the average SST in the 1990s, the 1960s were still relatively cool (Fig. 5a). It was also during the 1960s that underwater observations of BSB spawning aggregations were first documented (Fig. 7g). During this time, CDFW field biologists referred to BSB as “a more southern species frequenting our coast in and subsequent to periods of warmer waters,” and “Recently, 1960 to 1970, barred sand bass have formed an important part of the sport catch." (Fig. 7g, Feder et al. 1974).

Following the increase in Rockbass harvest during the 1960s, the Rockbass bag limit was increased in 1972 from 15 fish in combination with not more than ten of any one species, to 20 fish in combination with not more than ten of any one species (Fig. 7). Nevertheless, a year earlier, when reporting on the status of the BSB population, CDFW resource managers foreshadowed a decrease in BSB availability in southern California, “One cloud on the horizon—barred sand bass have not always been present in large numbers in southern California.” (Fig. 7h; Frey 1971). Shortly thereafter, harvest declined dramatically and the Rockbass bag limit was reduced by half to ten fish in combination. By the mid-to-late 1970s, Rockbass harvest returned to being dominated by Kelp Bass, and BSB CPFV CPUE was calculated to be 5-10x lower than was later observed in the 1980s during the subsequent warm regime (Fig. 7i, Love et al. 1996b).

**Table S1.** Prior parameter distributions used in the Bayesian capture-mark-reencounter models in this study. yal = years at liberty.

| Model | Parameter | θ | Distribution |
|---|---|---|---|
| 1960s and 1990s | true survival | $\phi$ | beta(1,1) |
| | recapture probability | $p$ | beta(1,1)* |
| | recovery probability | $\kappa$ | beta(1,1) |
| | resighting probability | $R$ | beta(1,1)* |
| | mean length at age 3 y | $L1$ | normal(236,10) |
| | mean length at age 9.5 y | $L2$ | normal(403,10) |
| | mean length at age 16 y | $L3$ | normal(495,10) |
| | probability tag retained after 1 yal | $tr_{1}$ | beta(1120,162) |
| | probability tag retained after 2 yal | $tr_{2}$ | beta(72,23) |
| | probability tag retained after 3 yal | $tr_{3}$ | beta(11,7) |
| | probability tag retained after 4 yal | $tr_{4}$ | beta(6,6) |
| | probability tag retained after 1 yal | $tr_{5}$ | beta(5,6) |
| | probability tag retained after 2 yal | $tr_{6}$ | beta(4,6) |
| | probability tag retained after 1 yal | $tr_{7}$ | beta(4,6) |
| | probability tag retained after 2 yal | $tr_{8}$ | beta(3,6) |
| | probability tag retained after 1 yal | $tr_{9}$ | beta(3,6) |
| | probability tag retained after 2 yal | $tr_{10}$ | beta(3,6) |
| | probability tag retained after 1 yal | $tr_{11}$ | beta(3,6) |
| 2010s | true survival | $\phi$ | beta(1,1) |
| | recapture probability | $p$ | beta(1,1) |
| | recovery probability | $\kappa$ | beta(1,1) |
| | resighting probability | $R$ | beta(1,1) |
| | mean length at age 24 mos | $L1$ | normal(191,100) |
| | mean length at age 108 mos | $L2$ | normal(391,100) |
| | mean length at age192 mos | $L3$ | normal(487,100) |
| | annual tag retention rate | $r$ | beta(140,27) |

*This parameter fixed at zero in the 1990s mark-resight-recovery model.

**Table S2.** Compilation of Barred Sand Bass (BSB) harvest statistics used in calculating the estimated mean annual numbers of legal- and sublegal-size fish harvested in the fishery during each tagging period. Prop. = proportion, CPFV = Commercial Passenger Fishing Vessel.

| | | Angler-intercept/Phone Survey Estimates[a] | | | | | | |
|---|---|---|---|---|---|---|---|---|
| Decade | Year | Prop. Legal Size | Total observed (measured) | Shore-based | Party/Charter Boats | Private/Rental Boats | Total BSB Harvest (all fishing modes) | CPFV BSB Harvest |
| 1960s[b] | 1964 | 0.85[c] | 5,562[c] | 7,318[b] | no estimate | 64,513[b] | 610,831 | 539,000[d] |
| | | | | | Prop. of Total | | | |
| | 1989 | 0.98 | 1,636 | -- | 0.59 | 0.40 | 1,295,773 | 787,074 |
| | 1993 | 0.97 | 2,086 | 0.00 | 0.56 | 0.43 | 731,182 | 309,000 |
| | 1994 | 0.97 | 1,393 | 0.03 | 0.54 | 0.43 | 703,763 | 270,000 |
| 1990s[e] | 1995 | 0.97 | -- | 0.02 | 0.64 | 0.34 | 801,512 | 349,000 |
| | 1996 | 0.97 | 1,948 | 0.01 | 0.68 | 0.32 | 743,805 | 591,000 |
| | 1997 | 0.98 | 1,062 | 0.02 | 0.41 | 0.57 | 462,973 | 476,000 |
| | 1998 | 0.98 | 1,460 | 0.01 | 0.37 | 0.62 | 417,633 | 376,000 |
| | 1999 | 0.98 | 3,925 | 0.00 | 0.44 | 0.56 | 488,743 | 414,000 |
| | 2013 | 0.91 | 1,031 | 0.05 | 0.62 | 0.34 | 64,796 | 56,000 |
| 2010s[f] | 2014 | 0.89 | 1,264 | 0.02 | 0.76 | 0.22 | 69,474 | 39,000 |

[a]Harvest estimates are provided for shore-based fishing (man-made structures, beach and bank) and boat-based fishing from CPFVs (Commercial Passenger Fishing Vessels; party/charter) and privately-owned/rental boats. Estimates are derived from a combination of angler intercept surveys and phone surveys of effort.

[b]Harvest estimates from Pinkas et al. (1968). The shore-based estimates are for the 1964/1965 season (July - June). The private boat estimates are for the year 1964 (January - December). Total harvest includes the estimate for CPFV

[c]Data source is for the year 1975, Wines (1978).

[d]Harvest estimates are for the year 1964 (January - December). Total bass harvested by CPFVs in 1964 was 1,078,000 fish; we applied a factor of 0.5 to this number to estimate BSB harvest; BSB comprised ~50% of the private boat harvest during this year (Pinkas et al. 1968).

[e]Survey proportions and harvest estimates obtained from the National Oceanic and Atmospheric Administration, Marine Recreational Fisheries Statistics Survey, 1980–2003.

[f]Survey proportions and harvest estimates obtained from the California Recreational Fisheries Survey, 2004–2021.

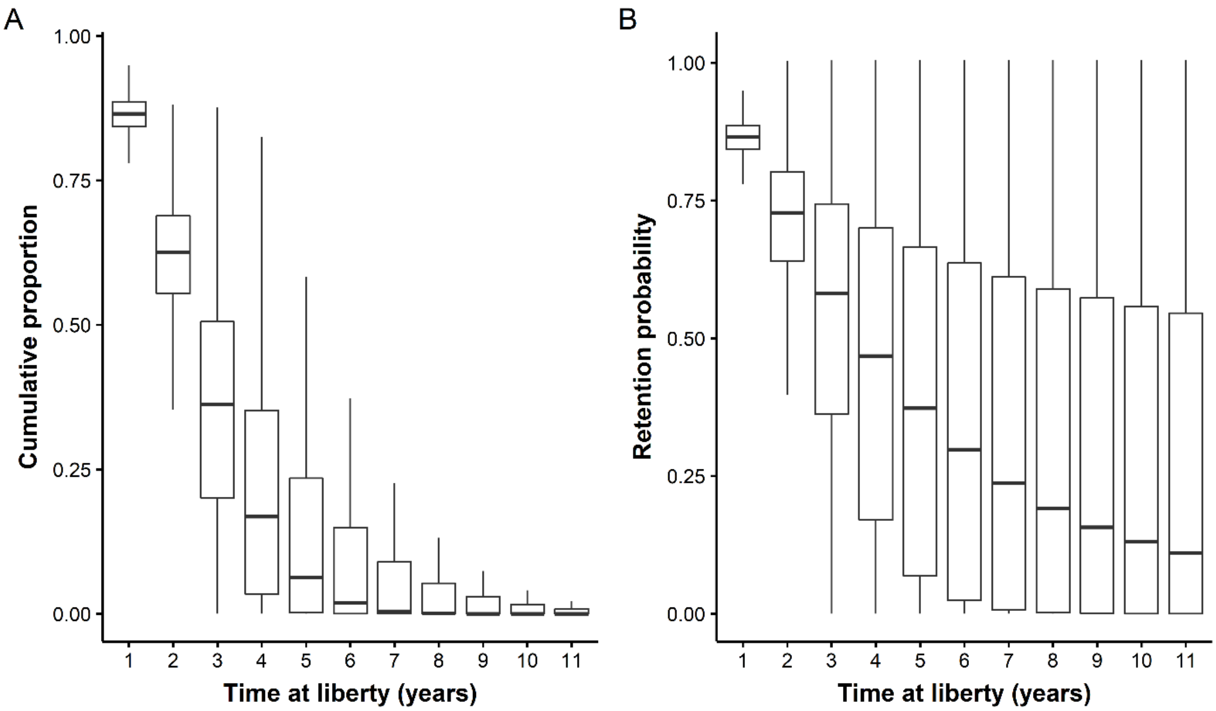


**Figure S1.** Box plots of Bayesian posterior estimates (mean and 95% Credible Intervals) of the a) cumulative proportion of double-tagged Kelp Bass retaining at least one tag over time, and b) the associated time-dependent tag retention probabilities applied as tag retention priors in the Barred Sand Bass capture-mark-reencounter models.

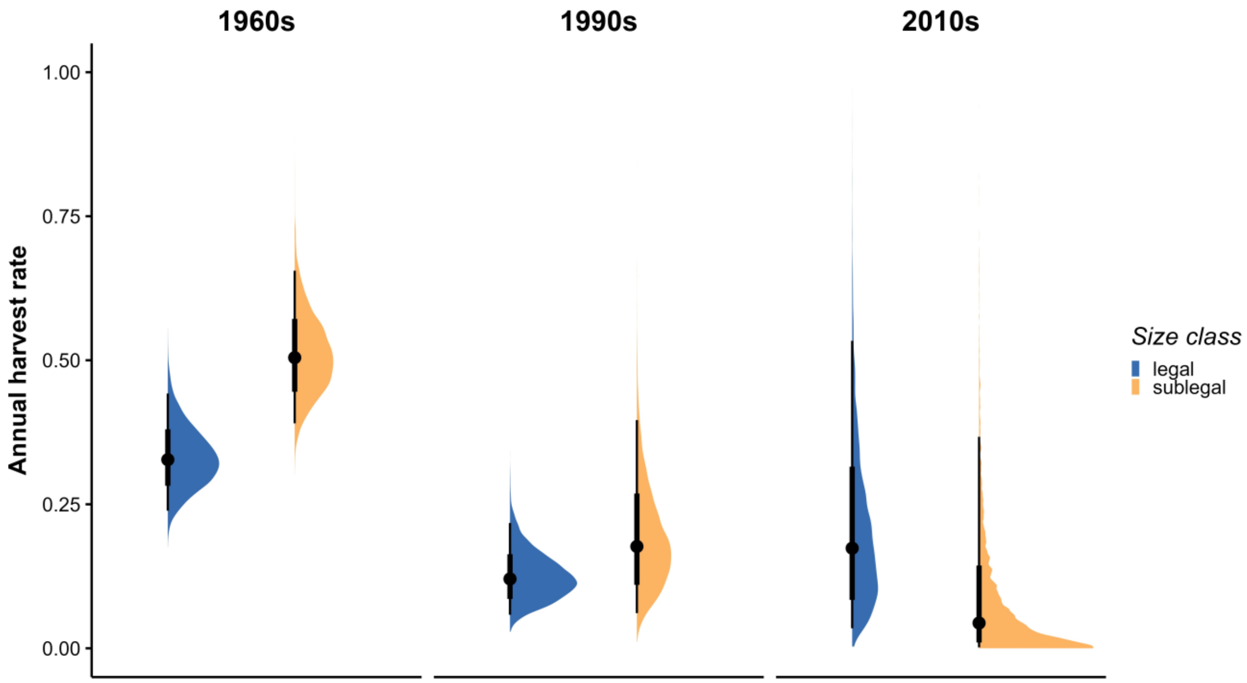


**Figure S2.** Bayesian capture-mark-reencounter model posterior distributions and mean and HDI 66 and 95% credible intervals (dots plus thick and thin lines) of annual harvest rates for legal- and sublegal-size Barred Sand Bass across tagging periods. The estimates are conservative, as they assume a 100% tag reporting rate.

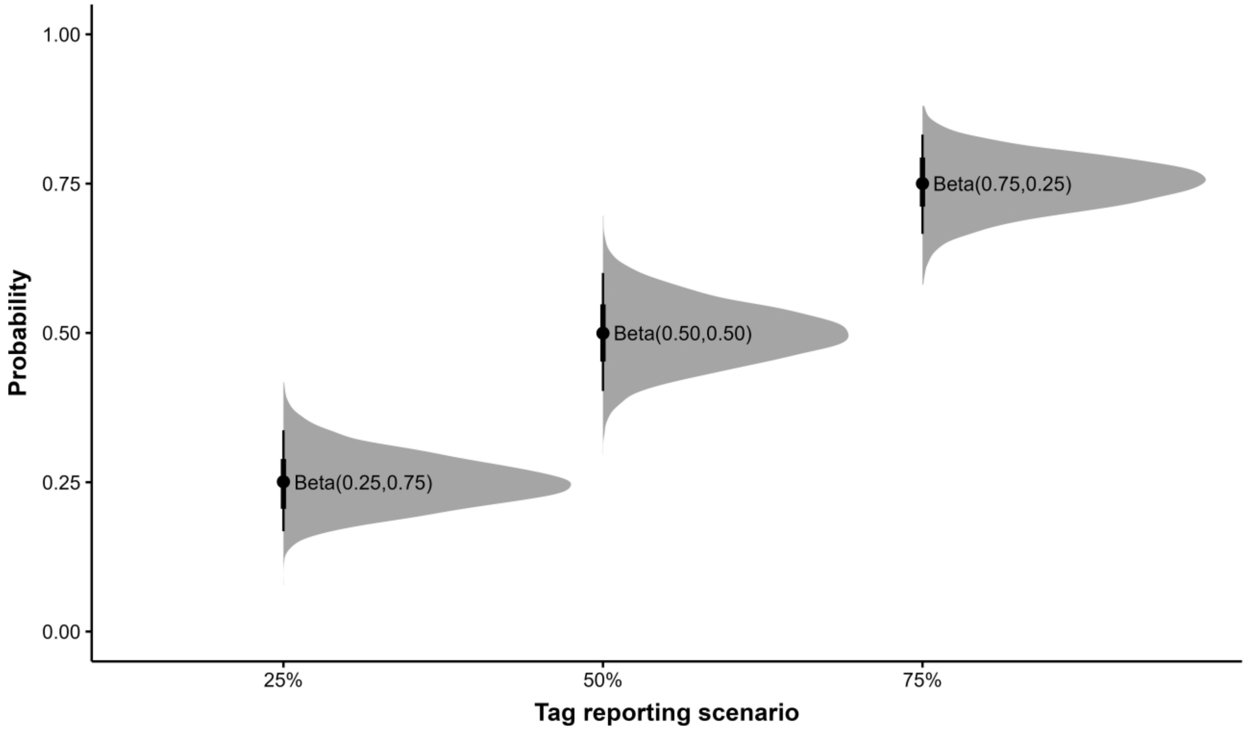


**Figure S3.** Bayesian prior distributions and mean and 66 and 95% HDI credible intervals (dots plus thick and thin lines) for the tag reporting prior sensitivity analysis used to derive harvest rates across three tag reporting rate scenarios. The assigned beta distributions are labeled for each scenario.

LITERATURE CITED


Bellquist LF (2015) A historical perspective of California recreational fisheries using a new database of 'trophy' fish records (1966-2013), combined with fisheries analyses of three species in the genus Paralabrax. PhD dissertation, University of California, San Diego, CA

CDFG [California Department of Fish and Game] (1962) Report for the month of August, 1962. Dept. of Fish and Game Records, Regional Reports, F3498:582-614.

Carlisle J, Turner C, Ebert E (1963) Artificial Habitat in the Marine Environment. Calif Dep Fish Game, Fish Bull 124.

Clark FN (1933) Rock bass (*Paralabrax*) in the California commercial fishery. Calif Fish Game 19:25–35.

Collyer R (1949) The Commercial Fish Catch of California for the Year 1947: With an Historical Review 1916-1947. Fish Bull:268

Croker RS (1940) Three Years of Fisheries Statistics on Marine Sport Fishing in California. Trans Am Fish Soc 69:111–118.

Feder HM, Turner CH, Limbaugh C (1974) Observations On Fishes Associated With Kelp Beds in southern California. Calif Dep Fish Game, Fish Bull 160

Frey HW (ed) (1971) Kelp and sand bass In: *California's Living Marine Resources and Their Utilization*. California Department of Fish and Game, Sacramento, USA, p 93-94.

Fry D, Croker RS (1934) A preliminary survey of Mission Bay State Park. Calif Fish Game 20:1–13.

Girard C (1858) Fishes. In: *General report upon the zoology of the several Pacific railroad routes. Explorations and surveys for a railroad route from the Mississippi River to the Pacific Ocean.* 10(4):1-400.

Holder CF (1912) The fishes of the Pacific coast, a handbook for sportsmen and tourists. Dodge Publishing Company, New York, USA.

Hubbs C (1948) Changes in the fish fauna of western north america correlated with changes in ocean temperature. J Mar Res VII: 459-482.

Jarvis ET, Gliniak HL, Valle CF (2014) Effects of fishing and the environment on the long-term sustainability of the recreational saltwater bass fishery in southern California. Calif Fish Game 100:234–259.

Love MS, Brooks A, Busatto D, Stephens J, Gregory PA (1996a) Aspects of the life histories of the kelp bass, *Paralabrax clathratus*, and barred sand bass, *P. nebulifer*, from the southern California Bight. Fish Bull 94:472–481.

Love MS, Brooks A, Ally JRR (1996b) An analysis of commercial passenger fishing vessel fisheries for kelp bass and barred sand bass in the southern California Bight. Calif Fish Game 82:105-121.

McClatchie S (2014) Regional Fisheries Oceanography of the California Current System. Springer Netherlands, Dordrecht.

Ogle DH, Doll J, Wheeler P (2022) Package ‘ FSA ’ version 0.9.3

Pinkas MS, Oliphant CW, Haugen L (1968) southern California marine sport-fishing survey: private boats, 1964; shoreline, 1965-66. Calif Dep Fish Game, Fish Bull 143

Plummer M, Stukalov A, Denwood M (2022) Package ‘ rjags ’ version 4-13.

Starks EC (1919) The basses and bass-like fishes of California. Families Serranidae, Haemulidae, Kyphosidae. Calif Fish Game 5:56–68.

Su Y-S, Yajima M (2021) Package ‘R2jags’ version 0.5-7

Turner CH, Ebert EE, Given RR (1969) Man-Made Reef Ecology. Calif Dep Fish Game, Fish Bull 146

Walker KM, Pentilla KM, Jarvis-Mason ET, Valle CF (2020) Validated age and growth of Barred Sand Bass within the Southern California Bight. Calif Fish Wildl J 106:205–220.

Wine V (1978) southern California Independent Sport Fishing Survey Annual Report No. 2. Mar Resour Adm Rep No 78-2:84.

Wine V (1979a) southern California Independent Sport Fishing Survey Annual Report No. 3. Mar Resour Adm Rep No 79-3:105.

Wine V (1979b) southern California Marine Sport Fishing: Private-Boat Catch and Effort, 1975-1976. Mar Resour Adm Rep No 79-11:64.

Young P (1969) The California Partyboat Fishery 1947–1967. Calif Dep Fish Game, Fish Bull 45

Young PH (1963) The kelp bass (*Paralabrax clathratus*) and its fishery, 1947-1958. Calif Dep Fish Game, Fish Bull 122
